# Modularity of genes involved in local adaptation to climate despite physical linkage

**DOI:** 10.1101/202481

**Authors:** Katie E. Lotterhos, Sam Yeaman, Jon Degner, Sally Aitken, Kathryn A. Hodgins

## Abstract

This preprint has been reviewed and recommended by Peer Community In Evolutionary Biology (https://doi.org/10.24072/pci.evolbiol.100050)

**Background:** Linkage among genes experiencing different selection pressures can make natural selection less efficient. Theory predicts that when local adaptation is driven by complex and non-covarying stresses, increased linkage is favoured for alleles with similar pleiotropic effects, with increased recombination favoured among alleles with contrasting pleiotropic effects. Here, we introduce a framework to test these predictions with a co-association network analysis, which clusters loci based on differing associations. We use this framework to study the genetic architecture of local adaptation to climate in lodgepole pine (*Pinus contorta*), based on associations with environments.

**Results:** We identified many clusters of candidate genes and SNPs associated with distinct environments (aspects of aridity, freezing, etc.), and discovered low recombination rates among some candidate genes in different clusters. Only a few genes contained SNPs with effects on more than one distinct aspect of climate. There was limited correspondence between co-association networks and gene regulatory networks. We further showed how associations with environmental principal components can lead to misinterpretation. Finally, simulations illustrated both benefits and caveats of co-association networks.

**Conclusions:** Our results supported the prediction that different selection pressures favored the evolution of distinct groups of genes, each associating with a different aspect of climate. But our results went against the prediction that loci experiencing different sources of selection would have high recombination among them. These results give new insight into evolutionary debates about the extent of modularity, pleiotropy, and linkage in the evolution of genetic architectures.

## Background

Pleiotropy and linkage are fundamental aspects of genetic architecture [1]. Pleiotropy is when a gene has effects on multiple distinct traits. Pleiotropy may hinder the rate of adaptation by increasing the likelihood that genetic changes have a deleterious effect on at least one trait [2, 3]. Similarly, linkage among genes experiencing different kinds of selection can facilitate or hinder adaptation [4–6]. Despite progress in understanding the underlying pleiotropic nature of phenotypes and the influence of pleiotropy on the rate of adaptation to specific conditions [7], we have an incomplete understanding of the extent and magnitude of linkage and pleiotropy in the local adaptation of natural populations to the landscapes and environments in which they are found.

Here, we aim to characterize the genetic architecture of adaptation to the environment, including the number of separate components of the environment in which a gene affects fitness (a form of “selectional pleiotropy,” Table 1)[8]. Genetic architecture is an encompassing term used to describe the pattern of genetic features that build and control a trait, and includes statements about the number of genes or alleles involved, their arrangement on chromosomes, the distribution of their effects, and patterns of pleiotropy (Table 1). We can measure many parameters to characterize environments *(e.g*., temperature, latitude, precipitation), but the variables we define may not correspond to the environmental factors that matter for an organism’s fitness. A major hurdle in understanding how environments shape fitness is defining the environment based on factors that drive selection and local adaptation, and not by the intrinsic attributes of the organism or by the environmental variables we happen to measure.

**Table 1.** Overview of terminology used in the literature regarding pleiotropy and modularity.

| Term | References | Meaning |
| --- | --- | --- |
| Genetic architecture | [1] | Genetic architecture refers to the pattern of genetic effects that build and control a facet of the organism (character, trait, or fitness). A description of genetic architecture includes statements about gene and allele number, the distribution of allelic and mutation effects, patterns of pleiotropy, and recombination rates among |
|  |  | causal loci on chromosomes. |
| Selectional pleiotropy | [8] | The number of separate components of fitness a mutation effects. Traits are defined by the action of selection and not by the intrinsic attributes of the organism. |
| Antagonistic pleiotropy at a single locus | [9] | In the context of this study, an allele exhibits antagonistic pleiotropy if it has different effects on fitness at different extremes of an environmental variable (e.g., positive effects on fitness in cold environments and negative effects in warm environments), which results in an association between the allele frequency and the environmental variable |
| Environmental pleiotropy | This study | Genes affect fitness in multiple distinct aspects of the multivariate environment, where each aspect is defined by the action of selection |
| Modularity or modular genetic architecture | [25] | A modular unit is a complex of elements (characters or genes) that: 1) collectively serve a similar functional role, 2) are tightly integrated by strong pleiotropic effects of genetic variation, and 3) are relatively independent from other such units. Pleiotropic effects may be on traits or on fitness, and are limited to elements within a module, with a suppression of pleiotropic effects between different modules (Figure 1A, left column). Genes within a module may or may not be physically linked. |
| Co-association network analysis | This study | An application of network theory used to identify modules of loci that are similar in their associations across many variables. |
| Co-association module | This study | A group of SNPs that show associations with a distinct *environmental factor*. These modules can be thought of as “variational” modules [sensu 19], which are composed of features that vary together and are relatively independent of other such sets of features. In practice, co-association modules are inferred by their similarity in associations with multiple environmental variables. |
| Selective environmental factors | This study | The specific aspect of the multivariate environment to which a SNP adapts on a geographic landscape. In practice, these are inferred by the environmental variables that associate with candidate SNPs within co-association modules. |

In local adaptation to climate, an allele that has different effects on fitness at different extremes of an environmental variable (e.g., positive effects on fitness in cold environments and negative effects in warm environments, often called “antagonistic pleiotropy”, Table 1[9]) will evolve to produce a clinal relationship between the allele frequency and that environmental factor [10–15]. While associations between allele frequencies and environmental factors have been well characterized across many taxa [16], whether genes affect fitness in multiple distinct aspects of the environment, which we call “environmental pleiotropy” (e.g., has effects on fitness in both cold and dry environments, Table 1), has not been well characterized [17]. This is because of conceptual issues that arise from defining environments along the univariate axes that we measure. For example, “cold” and “dry” might be a single selective optimum (“cold-dry”) to which a gene adapts [7], but these two axes are typically analyzed separately. Moreover, climate variables such as temperature and precipitation are highly correlated across landscapes, and this correlation structure makes inferring pleiotropy from signals of selection to climate difficult. Indeed, in their study of climate adaptation in *Arabidopsis*, Hancock et al. [17] noticed that candidate loci showed signals of selection in multiple environmental variables, potentially indicating pleiotropic effects. However, they also found that a substantial proportion of this overlap was due to correlations among climate variables on the landscape, and as a result they were unable to fully describe pleiotropic effects.

Because of the conceptual issues described above, certain aspects of the genetic architecture of adaptation to landscapes have not been well characterized, particularly the patterns of linkage among genes adapting to distinct environmental factors, and the degree of pleiotropic effects of genes on fitness in distinct environments. These aspects of genetic architecture are important to characterize in order to test the theoretical predictions described below, and to inform the considerable debate about whether organisms have a modular organization of gene effects on phenotypes or fitness components, versus universal effects of genes on all phenotypes or fitness components (Figure 1A, compare left to right column) [18–24].

**Figure 1.**
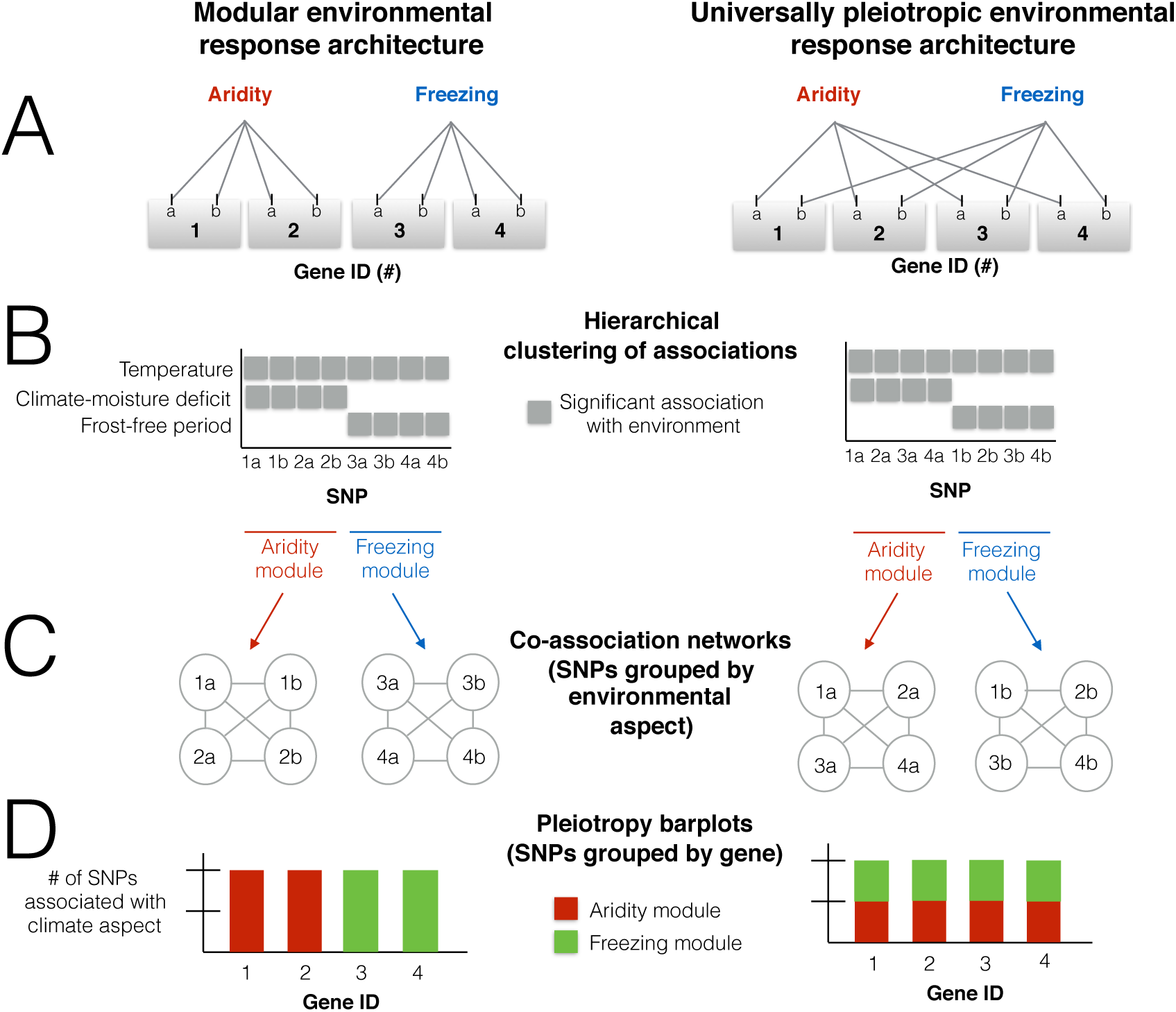
Conceptual framework for evaluating the modularity and pleiotropy of genetic architectures adapting to the environment. In this example, each gene (identified by numbers) contains two causal SNPs (identified by letters) where mutations affect fitness in potentially different aspects of the environment. The two aspects of the environment that affect fitness are aridity and freezing. A) The true underlying genetic architecture adapting to multiple aspects of climate. The left column represents a modular genetic architecture in which any pleiotropic effects of genes are limited to a particular aspect of the environment. The right column represents a non-modular architecture, in which genes have pleiotropic effects on multiple aspects of the environment. Universal pleiotropy occurs when a gene has effects on all the multiple distinct aspects of the environment. Genes in this example are unlinked in the genome, but linkage among genes is an important aspect of the environmental response architecture. B) Hierarchical clustering is used to identify the “co-association modules,” which jointly describe the groups of loci that adapt to a distinct aspects of climate as well as the distinct aspects of climate to which they adapt. In the left column, the “aridity module” is a group of SNPs within two unlinked genes adapting to aridity, and SNPs within these genes show associations with both temperature and climate-moisture deficit. In the right column, note how the aridity module is composed of SNPs from all 4 unlinked genes. C) Co-association networks are used to visualize the results of the hierarchical clustering with regards to the environment, and connections are based on similarity in SNPs in their associations with environments. In both columns, all SNPs within a module (network) all have similar associations with multiple environmental variables. D) Pleiotropy barplots are used to visualize the results of the hierarchical clustering with regards to the genetic architecture, represented by the proportion of SNPs in each candidate gene that affects different aspects of the environment (as defined by the co-association module).

Modular genetic architectures are characterized by extensive pleiotropic effects among elements within a module, and a suppression of pleiotropic effects between different modules [25]. Note that modularity in this study refers to similarity in the effects of loci on fitness, and not necessarily to the physical location of loci on chromosomes or to participation in the same gene regulatory network. Modular genetic architectures are predicted to be favored when genomes face complex spatial and temporal environments [26] or when multiple traits are under a combination of directional and stabilizing selection (because modularity allows adaptation to take place in one trait without undoing the adaptation achieved by another trait) [25, 27]. Adaptation to climate on a landscape fits these criteria because environmental variation among populations is complex - with multiple abiotic and biotic challenges occurring at different spatial scales - and traits are thought to be under stabilizing selection within populations but directional selection among populations [28].

Clusters of physically linked loci subject to the same selective environment, as well as a lack of physical linkage among loci subject to different selection pressures, are expected based on theory. When mutations are subject to the same selection pressure, recombination can bring variants with similar effects together and allow evolution to proceed faster [29]. Clusters of adaptive loci can also arise through genomic rearrangements that bring existing mutations together [30], or because new causal mutations linked to adaptive alleles have an increased establishment probability [31]. Similarly, clusters of locally adaptive loci are expected to evolve in regions of low recombination, such as inversions, because of the reduced gene flow these regions experience [32, 33]. In general, these linked clusters of adaptive loci are favored over evolutionary time because low recombination rates increase the rate at which they are inherited together. Conversely, selection will also act to disfavour linkage and increase recombination rates between genes adapting to different selection pressures [34–36]. Thus, genes adapting to different selection pressures would be unlikely to be physically linked or to have low recombination rates between them. In practice, issues can arise in inference because physical linkage will cause correlated responses to selection in neutral loci flanking a causal locus. Large regions of the genome can share similar patterns of association to a given environmental factor, such that many loci within a given candidate region are probably not causally responding to selection. Conversely, if linked genes are associated with completely different aspects of the selective environment, this is unlikely to arise by chance.

In summary, current analytical techniques have given limited insight into the genetic architectures of adaptation to environmental variation across natural landscapes. Characterizing the different aspects of the environment that act on genomes is difficult because measured variables are univariate and may not be representative of selection from the perspective of the organism, and because of spatial correlations among environmental variables. Even when many variables are summarized with ordination such as principal components, the axes that explain the most variation in physical environment don’t necessarily correspond to the axes that cause selection because the components are orthogonal [37]. Furthermore, the statistical methods widely used for inferring adaptation to climate are also univariate in the sense that they test for significant correlations between the frequency of a single allele and a single environmental variable [e.g., 38, 39, 40]. While some multivariate regression methods like redundancy analysis have been used to understand how multiple environmental factors shape genetic structure [41, 42], they still rely on ordination and have not been used to identify distinct evolutionary modules of loci.

Here, we aim fill this gap by presenting a framework for characterizing the genetic architecture of adaptation to the environment, through the joint inference of modules of loci that associate with distinct environmental factors that we call “co-association modules” (Table 1, Figure 1), as well as the distinct factors of the environment to which they associate. Using this framework, we can characterize some aspects of genetic architecture, including modularity and linkage, that have not been well studied in the adaptation of genomes to environments.

This framework is illustrated in Figure 1 for four hypothetical genes adapted to two distinct aspects of climate (freezing and aridity). In this figure we compare the patterns expected for (i) a modular architecture (left column, where pleiotropic fitness effects of a gene are limited to one particular climatic factor) to (ii) a highly environmentally pleiotropic architecture (right column, where genes have pleiotropic effects on adaptation to distinct climatic factors). Candidate SNPs are first identified by the significance of the univariate associations between allele frequency and the measured environmental variables, evaluated against what would be expected by neutrality. Then, hierarchical clustering of candidate-SNP allele associations with environments is used to identify co-association modules (Figure 1B) [43–45]. These modules can be visualized with a co-association network analysis, which identifies groups of loci that may covary with one environmental variable but covary in different ways with another, revealing patterns that are not evident through univariate analysis (Figure 1C). By defining the distinct aspects of the selectional environment (Table 1) for each module through their environmental associations, we can infer pleiotropic effects of genes through the associations their SNPs have with distinct selective environmental factors (Figure 1D). In this approach, the genetic effects of loci on different traits under selection are unknown, and we assume that each aspect of the multivariate environment selects for a trait or suite of traits that can be inferred by connecting candidate loci directly to the environmental factors that select for particular allelic combinations.

We apply this new approach to characterize the genetic architecture of local adaptation to climate in lodgepole pine (*Pinus contorta*) using a previously published exome capture dataset [46–48] from trees that inhabit a wide range of environments across their range, including freezing temperatures, precipitation, and aridity [49–52]. Lodgepole pine is a coniferous species inhabiting a wide range of environments in northwestern North America and exhibits isolation by distance population structure across the range [46]. Previous work based on reciprocal transplants and common garden experiments has shown extensive local adaptation [46, 53, 54]. We recently used this dataset to study convergent adaptation to freezing between lodgepole pine and the interior spruce complex (*Picea glauca* × *Picea engelmannii*) [46–48]. However, the comparative approach was limited to discovering parallel patterns between species, and did not examine selective factors unique to one species. As in most other systems, the genomic architecture in pine underlying local adaptation to the multivariate environment has not been well characterized, and our reanalysis yields several new biological insights overlooked by the comparative approach.

We evaluated the benefits and caveats of this new framework by comparing it with other multivariate approaches (based on principal components) and by evaluating it with simulated data. The evaluation with simulations yielded several important insights, including the importance of using strict criteria to exclude loci with false positive associations with environments. Thus, a key starting point for inferring co-association modules is a good set of candidate SNPs for adaptation. We developed this candidate set by first identifying top candidate genes for local adaptation (from a previously published set of genes that contained more outliers for genotype-environment associations and genotype-phenotype associations than expected by chance, [46]). We then identified top candidate SNPs within these top candidate genes as those whose allele frequencies were associated with at least one environmental variable above that expected by neutrality (using a criterion that excluded false positives in the simulated data described below). To this set of top candidate SNPs, we applied the framework outlined in Figure 1 to characterize environmental modularity and linkage of the genetic architecture. The power of our dataset comes from including a large number of populations inhabiting diverse environments (>250), the accurate characterization of climate for each individual with 22 environmental variables, a high-quality exome capture dataset representing more than 500,000 single-nucleotide polymorphisms (SNPs) in ∼29,000 genes [46–48], a mapping population that allows us to study recombination rates among genes, and an outgroup species that allowed us to determine the derived allele for most candidate SNPs. When such data is available, we find that this framework is useful for characterizing the environmental modularity and linkage relationships among candidate genes for local adaptation to multivariate environments.

## Results

### Top candidate genes and top candidates SNPs

The study of environmental pleiotropy and modularity is relevant only to loci under selection. In this study we identified a SNP as a top candidate based on whether (i) it was located within a top-candidate gene, and (ii) its allele frequency was associated with at least one environmental variable above and beyond what may be expected for neutrality. Our “top candidate” approach identified a total of 117 candidate genes out of a total of 29,920 genes. These contigs contained 801 top-candidate SNPs (out of 585,270 exome SNPs) that were strongly associated with at least one environmental variable and were likely either causal or tightly linked to a causal locus. This set of top candidate SNPs was enriched for *X^T^X* outliers (Supplemental Figure 1; *X^T^X* is an analog of *F_ST_* that measures differentiation in allele frequencies across populations). To elucidate patterns of multivariate association, we applied the framework described in Figure 1 to these 801 top candidate SNPs.

### Co-association modules

Hierarchical clustering and co-association network analysis of top candidate SNPs revealed a large number of co-association modules, each of which contains SNPs from one or more genes. Each co-association module is represented by one or more top candidate SNPs (represented by nodes) that are connected by edges. The edges are drawn between two SNPs if they have similar associations with the environment below a distance threshold. The distance threshold was determined by simulation as a number that enriched connections among selected loci adapting to the same environmental variable, and also decreased the number of connections to false positive loci (see *Results: Simulated datasets)*.

For the purposes of illustration, we classified SNPs into 4 main groups, each with several co-association modules, according to the kinds of environmental variables they were most strongly associated with: Aridity, Freezing, Geography, and an assorted group we bin as “Multi” (Figure 2A, B). Note that while we could have chosen a different number of groups, this would not have changed the underlying clustering of the SNPs revealed by co-association networks that is relevant to modularity (Figure 2B-F). This division of data into groups was necessary to produce coherent visual network plots and to make data analyses more computationally efficient (we found when there were more than ~20,000 edges in the data, computation and plotting of the network were not feasible with the package). Note that SNPs in different groups are more dissimilar to SNPs in other groups than to those in the same group (based on the threshold we used to determine edges) and would not be connected by edges in a co-association module. Interestingly, this clustering by association signatures does not closely parallel the correlation structure among environmental variables themselves. For example, continentality (TD), degree-days below 0 (DD_0), and latitude (LAT) are all relatively strongly correlated (> 0.5), but the “Freezing” SNPs are associated with continentality and degree-days below 0, but not latitude (Figure 2A, 2B).

**Figure 2.**
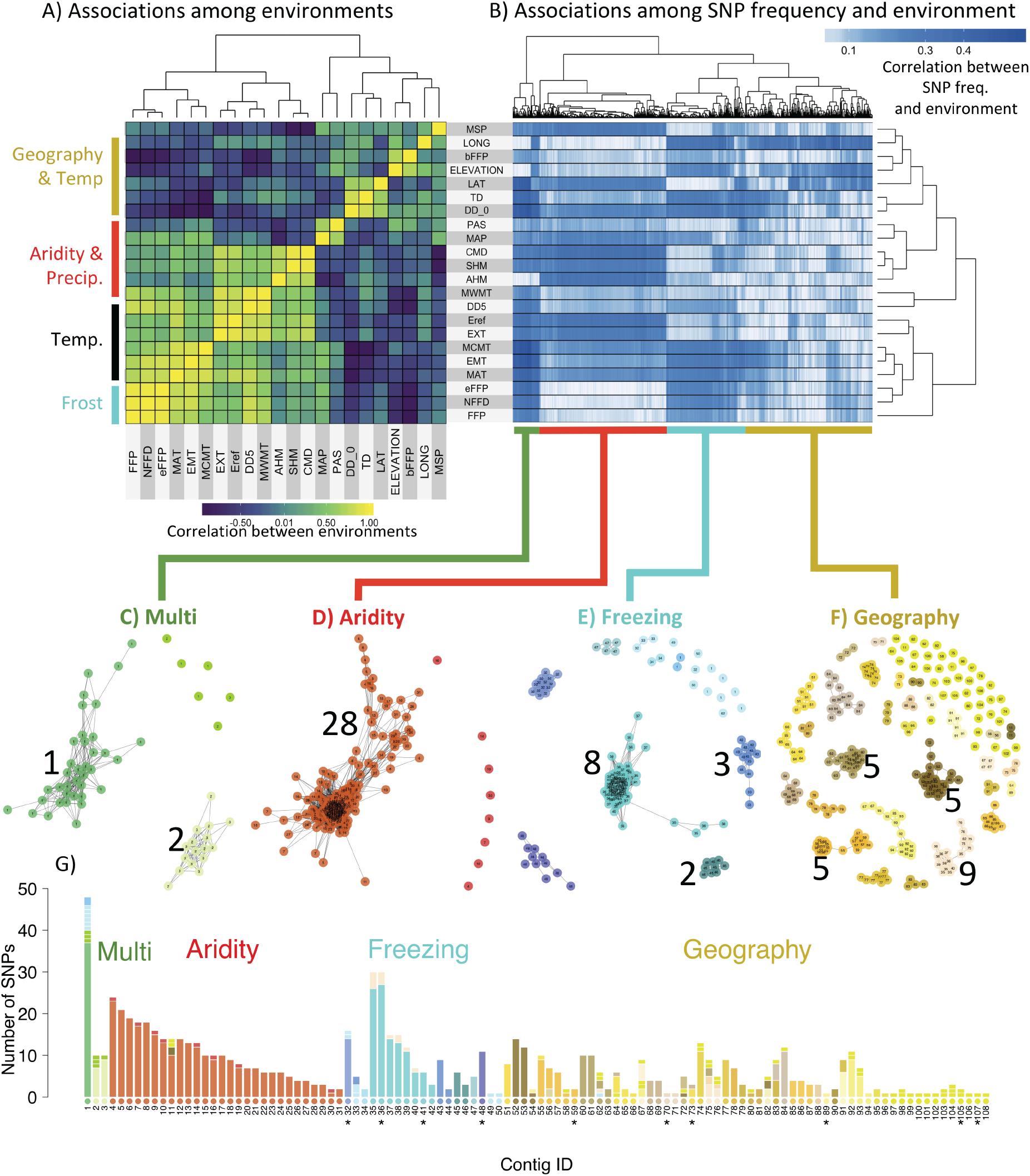
Co-association modules for *Pinus contorta*. A) Correlations among environments measured by Spearman’s ρ. Abbreviations of the environmental variables can be found in Table 2. B) Hierarchical clustering of associations between allele frequencies (of SNPs in columns) and environments (in rows) measured by Spearman’s ρ. C-F) Each co-association network represents a distinct co-association module, with color schemes according to the four major groups in the data. Each node is a SNP and is labeled with a number according to its exome contig, and a color according to its module - with the exceptions that modules containing a single SNP all give the same color within a major group. Numbers next to each module indicate the number of distinct genes involved (with the exception of the Geography group, where only modules with 5 or more genes are labeled). G) The pleiotropy barplot, where each bar corresponds to a contig, and the colors represent the proportion of SNPs in each co-association module. Note that contig IDs are ordered by their co-association module, and the color of contig-IDs along the x-axis is determined by the co-association module that the majority of SNPs in that contig cluster with. Contigs previously identified as undergoing convergent evolution with spruce by Yeaman et al. 2016 are indicated with “*”. Abbreviations: “Temp”: temperature, “Precip”: precipitation, “freq”: frequency.

**Table 2.**
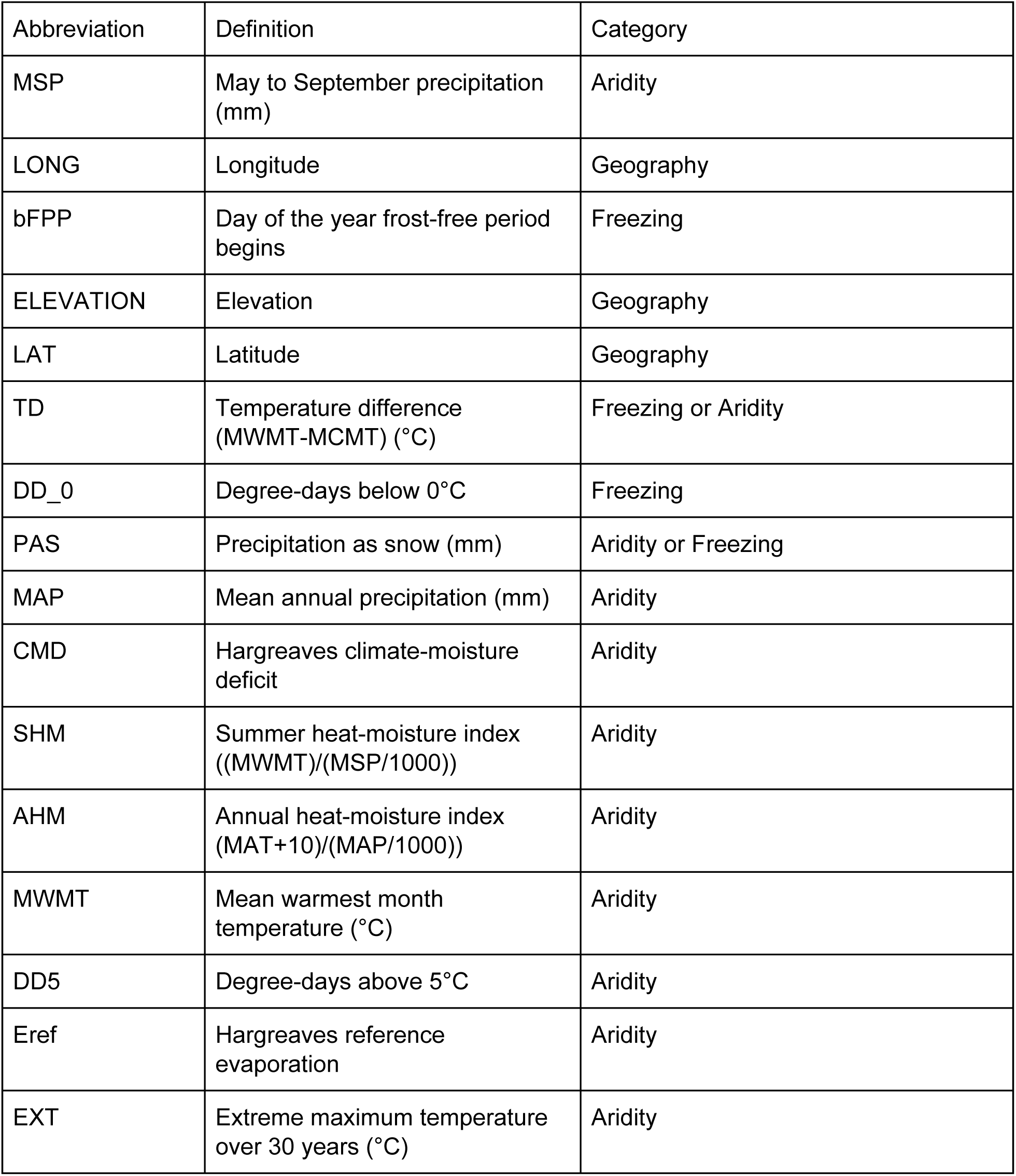
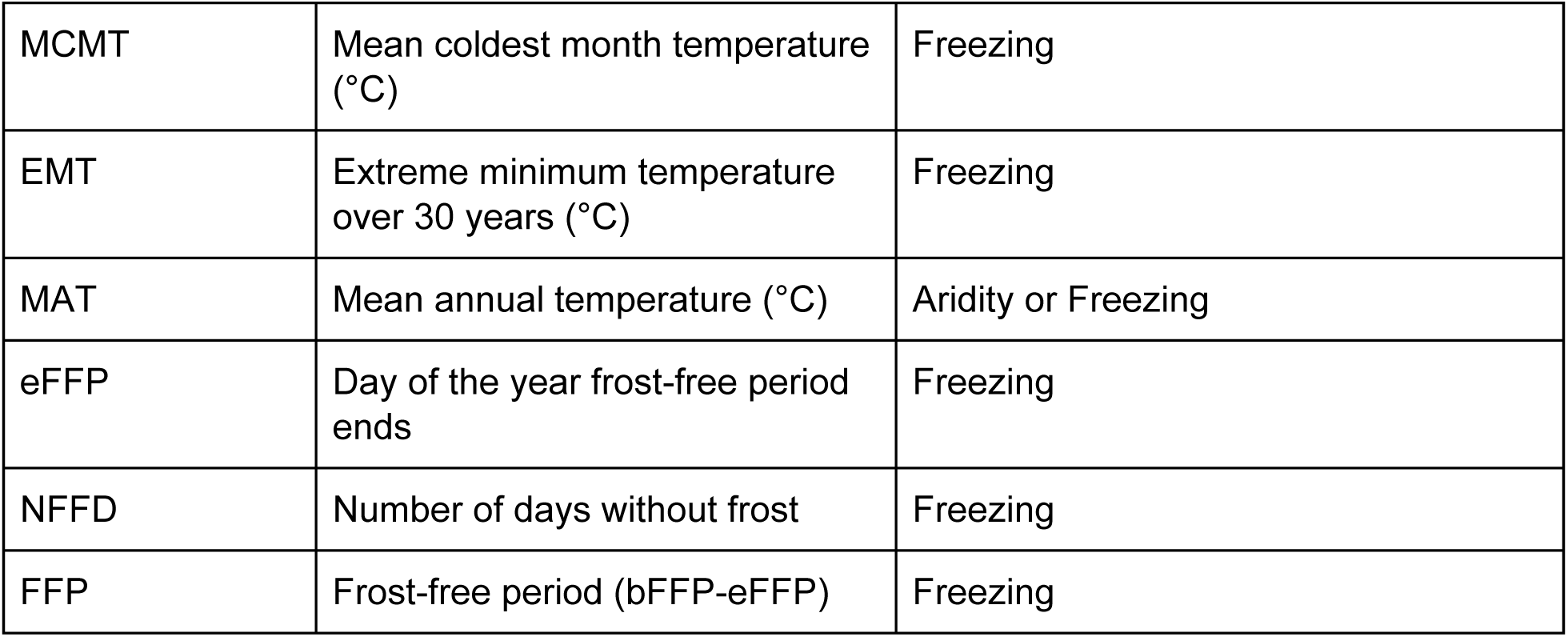
Environmental variables measured for each sampling location, ordered by their abbreviations shown in Figure 2 A and B.

The co-association modules are shown in Figures 2C-F. Each connected network of SNPs can be considered a group of loci that shows associations with a distinct environmental factor. The “Multi” group stands for multiple environments because these SNPs showed associations with 19 to 21 of the 22 environmental variables. This group consisted of 60 top candidate SNPs across just 3 genes, and undirected graph networks revealed 2 co-association modules within this group (Figure 2C, Supplementary Figure 2). The “Aridity” group consisted of 282 SNPs across 28 genes and showed associations with climate moisture deficit, annual heat:moisture index, mean summer precipitation, and temperature variables excluding those that were frost-related (Figure 2B). All these SNPs were very similar in their patterns of association and grouped into a single co-association module (Figure 2D, Supplementary Figure 3). The “Freezing” group consisted of 176 SNPs across 21 genes and showed associations with freezing variables including number of degree-days below 0°C, mean coldest month temperature, and variables related to frost occurrence (Figure 2B). SNPs from eight of the genes in this group formed a single module (genes #35–42), with the remaining SNPs mainly clustering by gene (Figure 2E, Supplementary Figure 4). The final group, “Geography,” consisted of 282 SNPs across 28 genes that showed consistent associations with the geographical variables elevation and longitude, but variable associations with other climate variables (Figure 2B). This group consisted of several co-association modules containing 1 to 9 genes (Figure 2F, Supplementary Figure 6). Network analysis using population-structure-corrected associations between allele frequency and the environmental variables resulted in broadly similar patterns, although the magnitude of the correlations was reduced (Supplemental Figure 6).

The pleiotropy barplot is visualized in Figure 2G, where each gene is listed along the x-axis, the bar color indicates the co-association module, and the bar height indicates the number of SNPs clustering with that module. If each co-association module associates with a distinct aspect of the multivariate environment, then genes whose SNPs associate with different co-association modules (e.g., genes with different colors in their bars in Figure 2G) might be considered to be environmentally pleiotropic. However, conceptual issues remain in inferring the extent of pleiotropy, because co-association modules within the Geography group, for instance, will be more similar to each other in their associations with environments than between a module in the Geography group and a module in the Multi group. For this reason, we are only inferring that our results are evidence of environmental pleiotropy when genes have SNPs in at least 2 of the 4 major groups in the data. For instance, gene #1, for which the majority of SNPs cluster with the Multi group, also has 8 SNPs that cluster with the Freezing group (although they are not located in co-association modules with any genes defined by Freezing). In the Aridity group, gene #11 has three SNPs that also cluster with the Geography group (although they are not located in co-association modules with any genes defined by Geography). In the Freezing group, some genes located within the same co-association module (#35–40) also have SNPs that cluster with another module in the Geography group (with genes #75–76; these are not physically linked to genes #35–37, see below). Whether or not these are “true” instances of environmental pleiotropy remains to be determined by experiments. For the most part, however, the large majority of SNPs located within genes are in the same co-association module, or in modules located within one of the four main groups, so environmental pleiotropy at the gene-level appears to be generally quite limited.

### Statistical and physical linkage disequilibrium

To determine if grouping of SNPs into co-association modules corresponded to associations driven by statistical associations among genes measured by linkage disequilibrium (LD), we calculated mean LD among all SNPs in the top candidate genes (as the correlation in allele frequencies). We found that the co-association modules captured patterns of LD among the genes through their common associations with environmental variables (Supplementary Figure S7). There was higher than average LD within the co-association modules of the Multi, Aridity, and Freezing groups, and very low LD between the Aridity group and the other groups (Supplementary Figure S7). The LD among the other three groups (Multi, Freezing, and Geography) was small, but higher with each other than with Aridity. Thus, the co-association clustering corresponded to what we would expect based on LD among genes, with the important additional benefit of linking LD clusters to likely environmental drivers of selection.

The high LD observed within the four main climate modules could arise via selection by the same factor of the multivariate environment, or via physical linkage on the chromosome, or both. We used a mapping population to disentangle these two hypotheses, by calculating recombination rates among the top candidate genes (see *Methods: Recombination rates*). Of the 117 top candidate genes, 66 had SNPs that were represented in our mapping population. The recombination data revealed that all the genes in the Aridity group have strong LD and are physically linked (Figure 3). Within the other three groups, we found physical proximity for only a few genes, typically within the same co-association module (but note that our mapping analysis does not have high power to infer recombination rate when loci are physically unlinked; see *Methods)*. For example, a few co-association modules in the Geography group (comprised of genes #53–54, #60–63, or #75–76) had very low recombination rates among them. Of the three genes forming the largest co-association module in the Freezing Group that was represented in our mapping panel (#35–37), two were physically linked.

**Figure 3.**
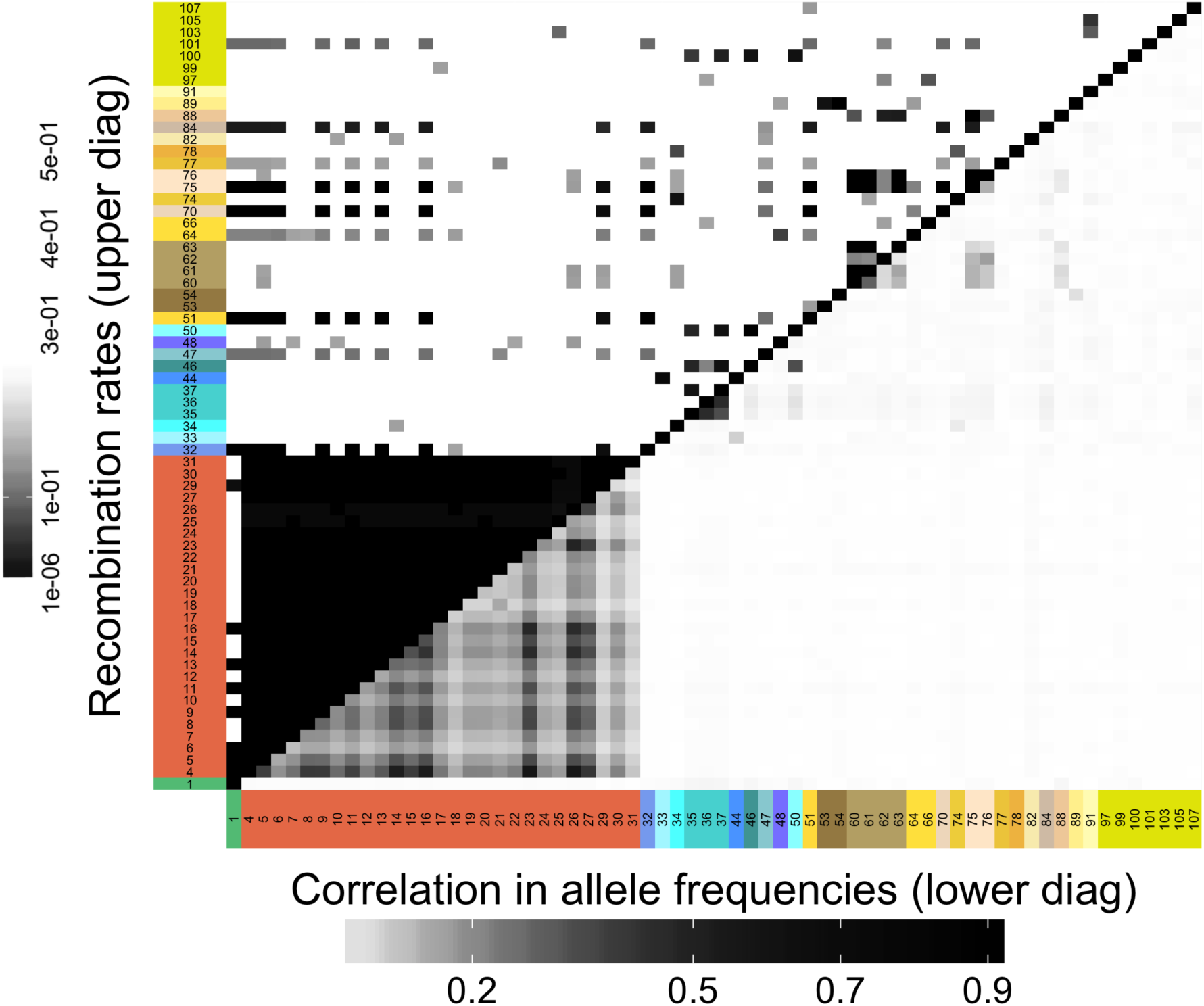
Comparison of linkage disequilibrium (lower diagonal) and recombination rates (upper diagonal) for exome contigs. Only contigs with SNPs in the mapping panel are shown. Rows and column labels correspond to Figure 2G. Darker areas represent either high physical linkage (low recombination) or high linkage disequilibrium.

Strikingly, low recombination rates were estimated between some genes belonging to different co-association modules across the four main groups, even though there was little LD among SNPs in these genes (Figure 3). This included a block of loci with low recombination comprised of genes from all 4 groups: 8 genes from the Aridity co-association module, 1 gene from the large module in the Multi group, 2 genes from different co-association modules in the Freezing group, and 7 genes from different co-association modules in the Geography group (upper diagonal of Figure 3, see Supplementary Figure S8 for a reorganization of the recombination data and more intuitive visualization).

### Comparison to conclusions based on principal components of environments

We compared the results from the co-association network analysis to associations with principal components (PC) of the environmental variables. Briefly, all environmental variables were input into a PC analysis, and associations between allele frequencies and PC axes were analyzed. We used the same criteria (log_10_ BF > 2 in bayenv2) to determine if a locus was a significant outlier and compared (i) overlap with top candidate SNPs based on outliers from univariate associations with environments, and (ii) interpretation of the selective environment based on loadings of environments to PC axes. The first three PC axes explained 44% (PC1), 22% (PC2), and 15% (PC3) of the variance in environments (80% total). Loadings of environment variables onto PC axes are shown in Supplementary Figure S9. A large proportion of top candidate SNPs in our study would not have been found if we had first done a PCA on the environments and then looked for outliers along PC axes: overall, 80% of the geography SNPs, 75% of the Freezing SNPs, 20% of the Aridity SNPs, and 10% of the Multi SNPs were *not* outliers along the first 10 PC axes and would have been missed.

Next, we evaluated whether interpretation of selective environments based on PCs was consistent with that based on associations with individual environmental factors. Some of the temperature and frost variables (MAT: mean annual temperature, EMT: extreme minimum temperature, DD0: degree days below 0C, DD5: degree days above 5C, bFFP: begin frost-free period, FFP: frost free period, eFFP: end frost free period, labels in Figure 2A) had the highest loadings for PC1 (Supplementary Figure S9). Almost all of the SNPs in the Multi group (90%) and 19% of SNPs in the Freezing group were outliers along this axis (Supplementary Figure 10, note green outliers along x-axis from Multi group; less than 2% of candidate SNPs in the other groups were outliers). For PC1, interpretation of the selective environment (e.g., MAT, DD0, FFP, eFFP, DD5) is somewhat consistent with the co-association network analysis (both Multi SNPs and Freezing SNPs show associations with all these variables, Figure 2B). However, the Multi SNPs and Freezing SNPs had strong associations with other variables (e.g., Multi SNPs showed strong associations with Latitude and Freezing SNPs showed strong associations with longitude, Figure 2B) that did not load strongly onto this axis, and would have been missed in an interpretation based on associations with principal components.

Many precipitation and aridity variables loaded strongly onto PC2, including mean annual precipitation, annual heat:moisture index, climate moisture deficit, and precipitation as snow (Supplementary Figure 9). However, few top candidate SNPs were outliers along the PC2 axis: only 13% of Freezing SNPs, 10% of Aridity SNPs, and less than 3% of Multi or Geography SNPs were outliers (Supplementary Figure 10A, note lack of outliers on y-axis).

For PC3, latitude, elevation, and two frost variables (beginning frost-free period and frost-free period) had the highest loadings (Supplementary Figure 9). The majority (78%) of the Aridity SNPs were outliers with PC3 (Supplementary Figure 10B, note outliers as orange dots on y-axis). Based on the PC association, this would lead one to conclude that the Aridity SNPs show associations with latitude, elevation, and frost-free period. While the Aridity SNPs do have strong associations with latitude (5th row in Figure 2B), they show very weak associations with the beginning of frost-free period, elevation, and frost-free period length (3rd, 4th, and last row in Figure 2B, respectively). Thus, interpretation of the environmental drivers of selection based on associations with PC3 would have been very different from the univariate associations.

### Interpretation of multivariate allele associations

While the network visualization gave insight into patterns of LD among loci, it does not give insight into patterns of allele frequency change on the landscape, relative to the ancestral state. As illustrated above, principal components would not be useful for this latter visualization. Instead, we accomplished this by plotting the association of a derived allele with one environmental variable against the association of that allele with a second environmental variable. Note that when the two environmental variables themselves are correlated on the landscape, an allele with a larger association in one environment will also have a larger association with a second environment, regardless of whether or not selection is shaping those associations. We can visualize (i) the expected genome-wide covariance (given correlations between environmental variables; Fig 1A left panel) using shading of quadrants and (ii) the observed genome-wide covariance using a 95% prediction ellipse (Figure 4). Since alleles were coded according to their putative ancestral state in loblolly pine *(Pinus taeda)*, the location of any particular SNP in the plot represents the bivariate environment in which the derived allele is found in higher frequency than the ancestral allele (Figure 4). Visualizing the data in this way allows us to understand the underlying correlation structure of the data, as well as to develop testable hypotheses about the true selective environment and the fitness of the derived allele relative to the ancestral allele.

**Figure 4.**
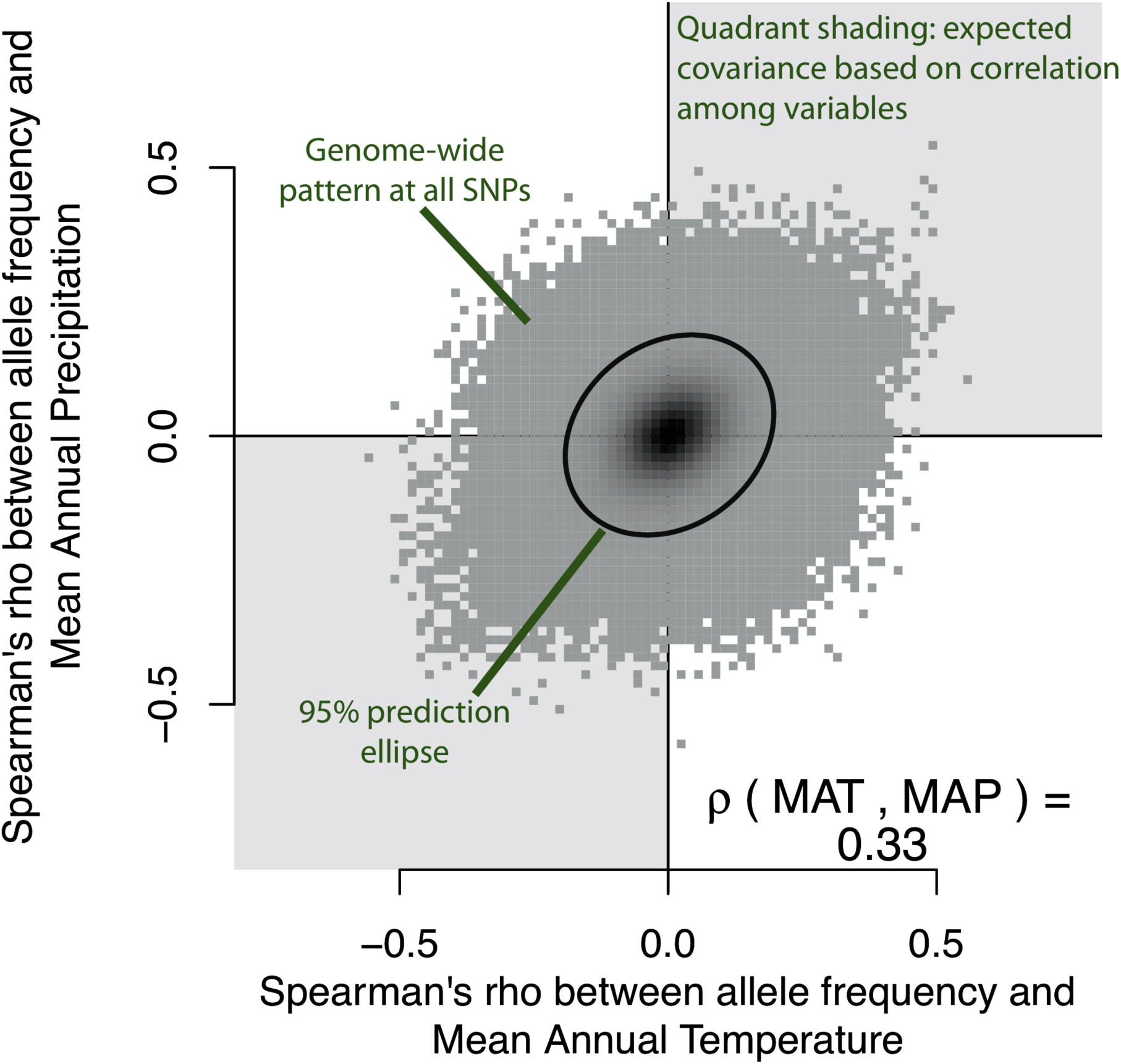
Overview of galaxy biplots. The association between allele frequency and one variable is plotted against the association between allele frequency and a second variable. The Spearmans? correlation between the two variables (mean annual temperature or MAT and mean annual precipitation or MAP in this example) is shown in the lower right corner. When the two variables are correlated, genome-wide covariance is expected to occur in the direction of their association (shown with quadrant shading in light grey). The observed genome-wide distribution of allelic effects is plotted in dark grey and the 95% prediction ellipse is plotted as a black line. Because derived alleles were coded as 1 and ancestral alleles were coded as 0, the location of any particular SNP in bivariate space represents the type of environment that the derived allele is found in higher frequency, whereas the location of the ancestral allele would be a reflection through the origin (note only derived alleles are plotted).

We overlaid the top candidate SNPs, colored according to their grouping in the co-association network analysis, on top of this genome-wide pattern (for the 668 of 801 top candidates for which the derived allele could be determined). We call these plots “galaxy biplots” because of the characteristic patterns we observed when visualizing data this way (Figure 5). Galaxy biplots revealed that SNPs in the Aridity group showed associations with hot/dry versus cold/wet environments (red points in Figure 5A), while SNPs in the Multi and Freezing groups showed patterns of associations with hot/wet versus cold/dry environments (blue and green dots in Figure 5A). These outlier patterns became visually stronger for some SNPs and environments after correcting associations for population structure (compare Figure 5A to Figure 5B, structure-corrected allele frequencies calculated with Bayenv2, see *Methods)*. Most SNPs in the Freezing group showed associations with elevation but not latitude (compare height of blue points on y-axis of Figure 5C to Figure 5E). Conversely, the large co-association module in the Multi group (gene #1, dark green points) showed associations with latitude but not elevation, while the second co-association module in the Multi group (genes #2–3, light green points) showed associations with both latitude and elevation (compare height of points on y-axis of Figure 5C to Figure 5E). Note how the structure correction polarized these patterns somewhat without changing interpretation, suggesting that the structure-corrected allelic associations become more extreme when their pattern of allele frequency contrasted the background population structure (compare left column of Figure 5 to right column of Figure 5).

**Figure 5.**
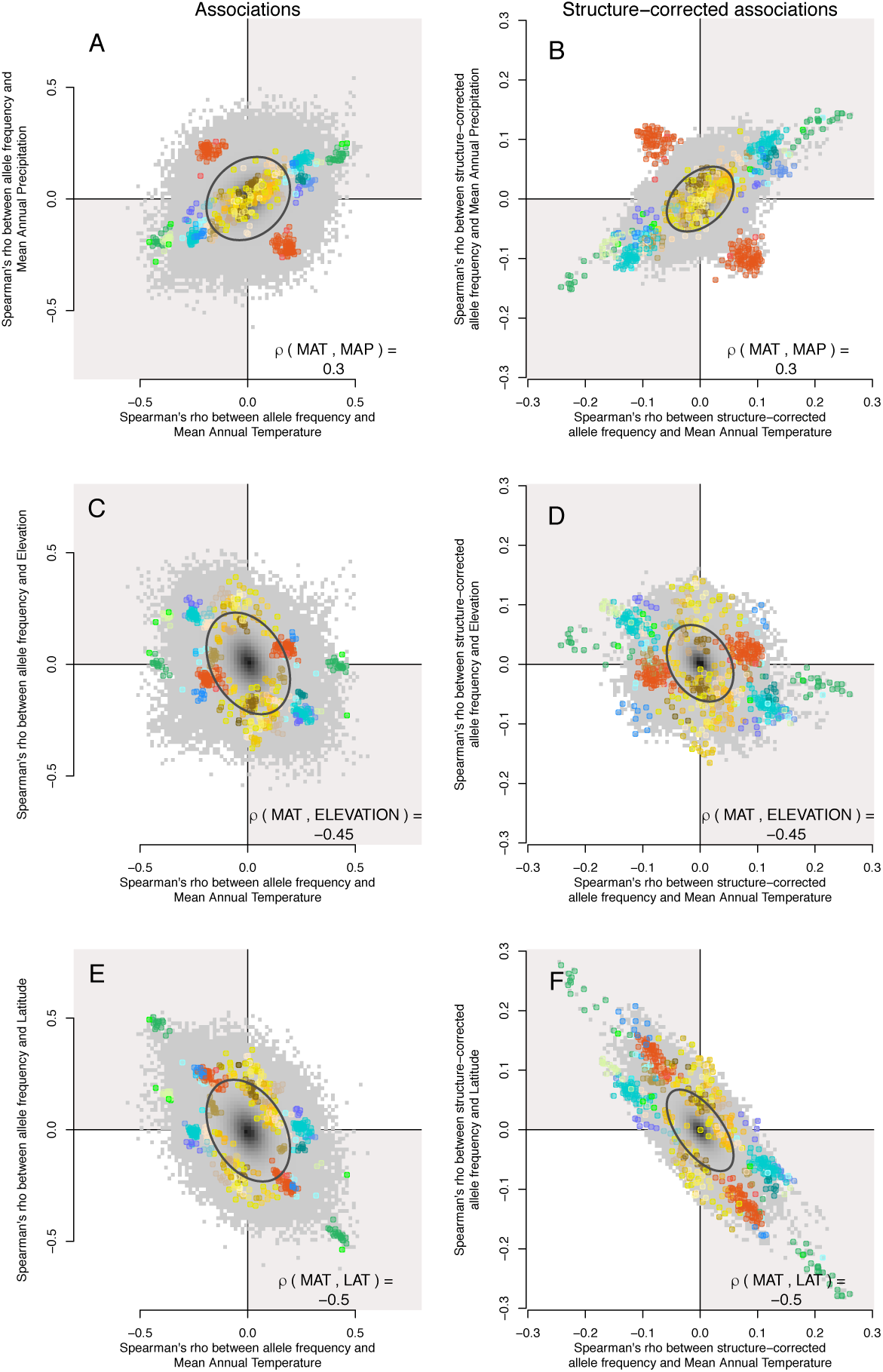
Galaxy biplots for different environmental variables for regular (left column) and structure-corrected (right column) associations. Top candidate SNPs are highlighted against the genome-wide background. The internal color of each point corresponds to its co-association module (as shown in Figure 2 C-F). Top row: mean annual temperature (MAT) vs. mean annual precipitation (MAP), middle row: MAT and Elevation, bottom row: MAT and latitude (LAT).

Some modules were particularly defined by the fact that almost all the derived alleles changed frequency in the same direction (e.g., sweep-like signatures). For instance, for the co-association module in the Multi group defined by genes #2–3, 14 of the 16 derived SNPs were found in higher frequencies at colder temperatures, higher elevations, and higher latitudes. Contrast this with a group of SNPs from an co-association module in the Freezing group defined by gene #32, in which 14 of 15 derived SNPs were found in higher frequencies in warmer temperatures and lower elevations, but showed no associations with latitude. These may be candidates for genotypes that have risen in frequency to adapt to particular environmental conditions on the landscape.

Conversely, other modules showed different combinations of derived alleles that arose in frequency at opposite values of environmental variables. For instance, derived alleles in the Aridity co-association module were found in higher frequency in either warm, dry environments (88 of 155 SNPs) or in cold, moist environments (67 of 155 SNPs). Similarly, for the Multi co-association module defined by gene #1, derived alleles were found in higher frequency in either cold, dry environments (15 of 37 SNPs) or in warm, moist environments (22 of 37 SNPs). These may be candidates for genes acted on by antagonistic pleiotropy within a locus (Table 1), in which one genotype is selected for at one extreme of the environment and another genotype is selected for at the other extreme of the environment. Unfortunately, we were unable to fully characterize the relative abundance of sweep-like vs. antagonistically pleiotropic patterns across all top candidate genes due to (i) the low number of candidate SNPs for most genes, and (ii) for many SNPs the derived allele could not be determined (because there was a missing SNP or other missing data in the ancestral species).

We also visualized the patterns of allele frequency on the landscape for two representative SNPs, chosen because they had the highest number of connections in their co-association module (and were more likely to be true positives, see *Results: Simulated datasets)*. Geographic and climatic patterns are illustrated with maps for two such SNPs: (i) a SNP in the Multi co-association module defined by gene #1 is shown in Figure 6A (with significant associations with latitude and mean annual temperature), and (ii) a SNP in the Aridity co-association module (Figure 6B, gene #8 from Figure 2, with significant associations with annual heat:moisture index and latitude). These maps illustrate the complex environments that may be selecting for particular combinations of genotypes despite potentially high gene flow in this widespread species.

### Candidate gene annotations

Although many of the candidate genes were not annotated, as is typical for conifers, the genes underlying adaptation to these environmental gradients had diverse putative functions. The top candidate SNPs were found in 3’ and 5’ untranslated regions and open reading frames in higher proportions than all exome SNPs (Supplemental Figure S11). A gene ontology (GO) analysis using previously assigned gene annotations [46, 55] found that a single molecular function, solute:cation antiporter activity, was over-represented across all top candidate genes (Supplemental Table S1). In the Aridity and Geography groups, annotated genes included sodium or potassium ion antiporters (one in Aridity, a KEA4 homolog, and two in Geography, NHX8 and SOS1 homologs), suggestive of a role in drought, salt or freezing tolerance [56]. Genes putatively involved in auxin biosynthesis were also identified in the Aridity (YUCCA 3) and Geography (Anthranilate synthase component) groups (Supplemental Table S2), suggestive of a role in plant growth. In the Freezing and Geography groups, several flowering time genes were identified [57] including a homolog of CONSTANS [58] in the Freezing group and a homolog of FY, which affects FCA mRNA processing, in the Geography group [58] (Supp Table 2). In addition, several putative drought/stress response genes were identified, such as DREB transcription factor [59] and an RCD1-like gene (Supplemental Table 2). RCD-1 is implicated in hormonal signaling and in the regulation of several stress-responsive genes in *Arabidopsis thaliana* [57]. In the Multi group, the only gene that was annotated functions in acclimation of photosynthesis to the environment in *A. thaliana* [60].

**Figure 6.**
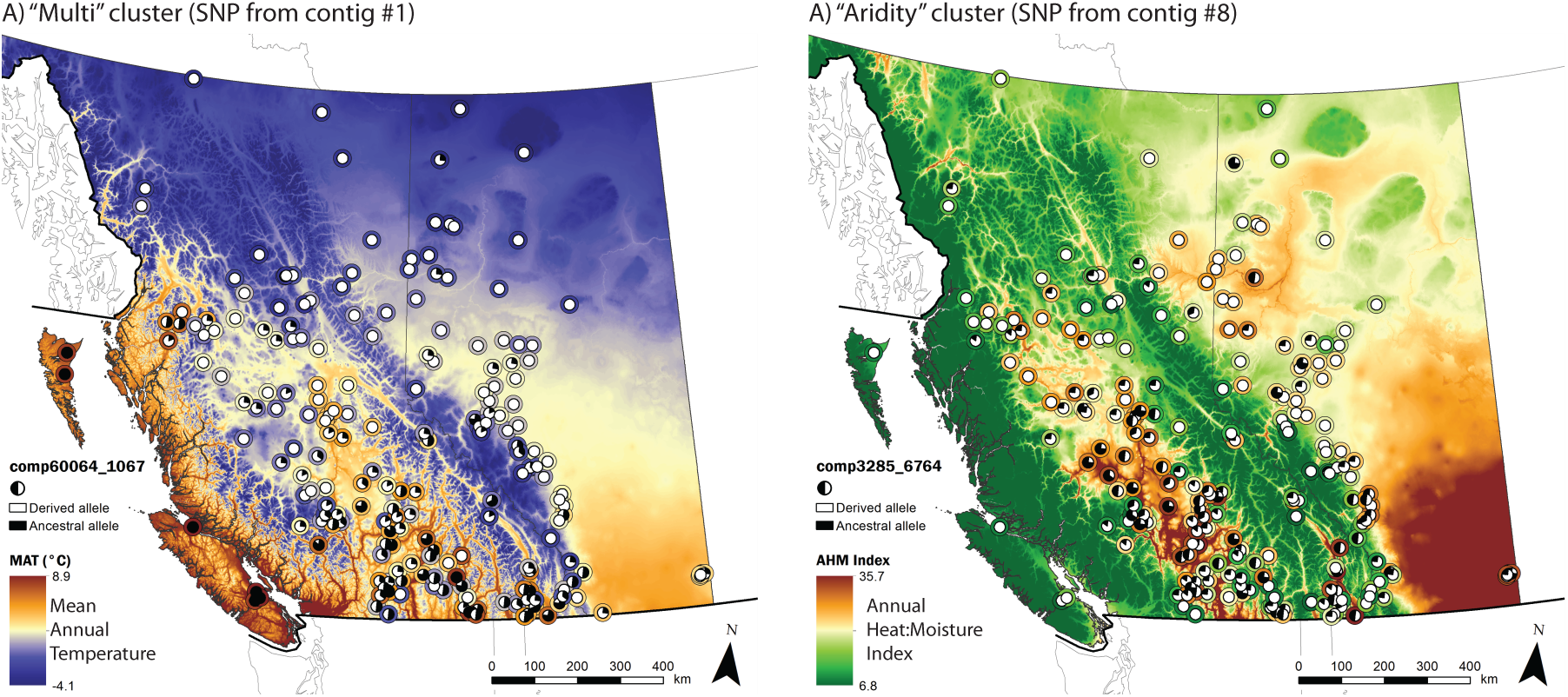
Pie charts representing the frequency of derived candidate alleles across the landscape. Allele frequency pie charts are overlain on top of an environment that the SNP shows significant associations with. The mean environment for each population is shown by the color of the outline around the pie chart. A) Allele frequency pattern for a SNP from contig 1 in the Multi cluster from Figure 2. The derived allele had negative associations with temperature but positive associations with latitude. B) Allele frequency pattern for a SNP from contig 8 in the Aridity cluster. The derived allele had negative associations with annual:heat moisture index (and other measures of aridity) and positive associations with latitude. SNPs were chosen as those with the highest degree in their co-association module.

Of the 47 candidate genes identified by Yeaman et al. 2016 as undergoing convergent evolution for adaptation to low temperatures in lodgepole pine and the interior spruce hybrid complex *(Picea glauca, P. engelmannii*, and their hybrids), 10 were retained with our stringent criteria for top candidates. All of these genes grouped into the Freezing and Geography groups (shown by “*” in Figure 2G): the two groups that had many SNPs with significant associations with elevation. This is consistent with the pattern of local adaptation in the interior spruce hybrid zone, whereby Engelmann spruce is adapted to higher elevations and white spruce is adapted to lower elevations [61].

### Comparison of co-expression clusters to co-association modules

To further explore if co-association modules have similar gene functions, we examined their gene expression patterns in response to climate treatments using previously published RNAseq data of 10,714 differentially expressed genes that formed 8 distinct co-expression clusters [55]. Of the 108 top candidate genes, 48 (44%) were also differentially expressed among treatments in response to factorial combinations of temperature (cold, mild, or hot), moisture (wet vs. dry), and/or day length (short vs. long day length). We found limited correspondence between co-association modules and co-expression clusters. Most of the top-candidate genes that were differentially expressed mapped to 2 of the 10 co-expression clusters previously characterized by [55] (Figure 7, blue circles are the P2 co-expression cluster and green triangles are the P7 co-expression cluster previously described by [55]). Genes in the P2 co-expression cluster had functions associated with the regulation of transcription and their expression was strongly influenced by all treatments, while genes in the P7 co-expression cluster had functions relating to metabolism, photosynthesis, and response to stimulus [55]. Genes from the closely linked Aridity group mapped to 4 distinct co-expression clusters, contigs from the Freezing group mapped to 3 distinct co-expression clusters, and genes from the Geography group mapped to 3 distinct co-expression clusters.

We used a Fisher exact test to determine if any co-expression cluster was over-represented in any of the the four major co-association groups shown in Figure 2. We found that the Freezing group was over-represented in the P2 co-regulated gene expression cluster *(P* < 0.05) with seven (58%) of the Freezing genes found within the P2 expression cluster, revealing coordinated expression in response to climatic conditions. Homologs of four of the seven genes were present in *A. thaliana*, and three of these genes were transcription factors involved in abiotic stress response *(DREB* transcription factor), flowering time (*CONSTANS*, pseudoresponse regulator) or the circadian clock (pseudo-response regulator 9). No other significant over-representation of gene expression class was identified for the four association groups or for all adaptation candidate genes.

**Figure 7.**
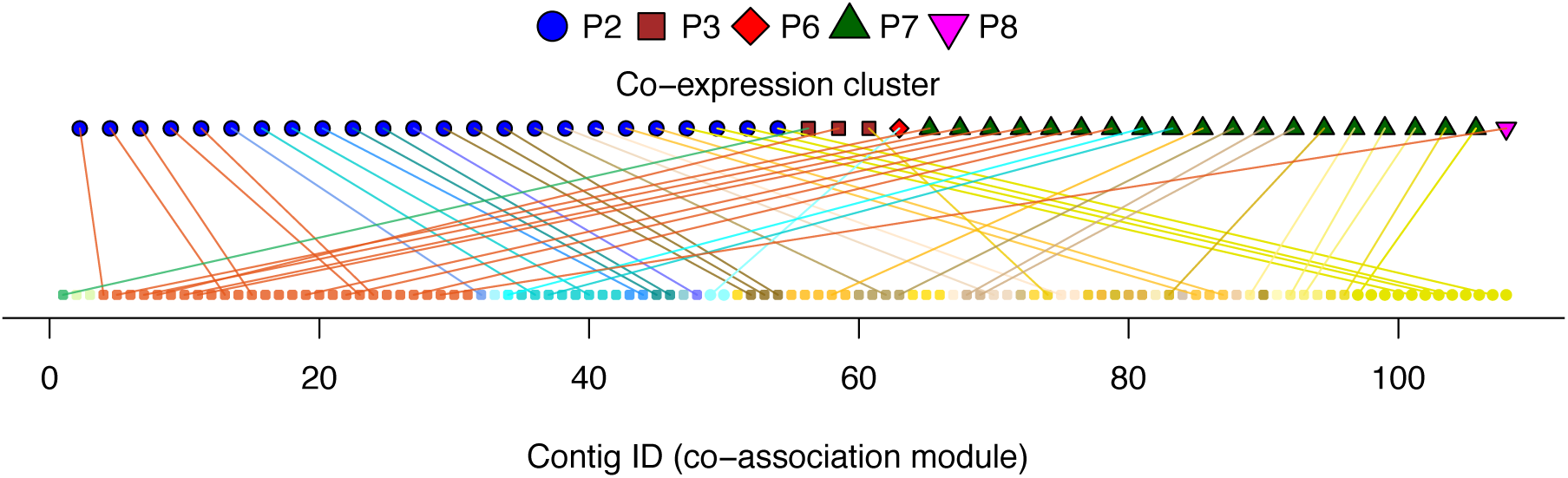
Co-association modules mapped to co-expression clusters determined by climate treatments. Contig ID, color, and order shown on the bottom correspond to co-association modules plotted in Figure 2. Co-expression clusters from [55] are shown at the top.

### Simulated datasets

We used individual-based simulations to examine potential limitations of the co-association network analysis by comparing the connectedness of co-association networks arising from false positive neutral loci vs. a combination of false positive neutral loci and true positive loci that had experienced selection to an unmeasured environmental factor. Specifically, we used simulations with random sampling designs from three replicates across three demographic histories: (i) isolation by distance at equilibrium (IBD), and non-equilibrium range expansion from a (ii) single refuge (1R) or from (iii) two refugia (2R). These landscape simulations were similar to lodgepole pine in the sense that they simulated large effective population sizes and resulted in similar *F_ST_* across the landscape as that observed in pine ([62, 63], *F_ST_* in simulations ~ 0.05, vs. F_ST_ in pine ~ 0.016 [46]). To explore how the allele frequencies that evolved in these simulations might yield spurious patterns under the co-association network analysis, we overlaid the 22 environmental variables used in the lodgepole pine dataset onto the landscape genomic simulations [62, 63]. To simulate selection to an unmeasured environmental factor, a small proportion of SNPs (1%) were subjected to computer-generated spatially varying selection along a weak latitudinal cline [62, 63]. We assumed that 22 environmental variables were measured, but not the “true” selective environment; our analysis thus represents the ability of co-association networks to correctly cluster selected loci even when the true selective environment was unmeasured, but a number of other environmental variables were measured (correlations between the selective environment and the other variables ranged from 0 to 0.2). Note that the simulations differ from the empirical data in at least two ways: (i) there is only one selective environment (so we can evaluate whether a single selective environment could result in multiple co-association modules in the data given the correlation structure of observed environments), and (ii) loci were unlinked.

The *P*-value and Bayes factor criteria for choosing top candidate SNPs in the empirical data produced no false positives with the simulated datasets (Supplemental Figure 12 right column), although using these criteria also reduced the proportion of true positives. Therefore, we used less stringent criteria to analyze the simulations so that we could also better understand patterns created by unlinked, false positive neutral loci (Supplemental Figure 12 left column).

We found that loci under selection by the same environmental factor generally formed a single tightly connected co-association module even though they were unlinked, and that the degree of connectedness of selected loci was greater than among neutral loci (Figure 8). Thus, a single co-association module typically resulted from adaptation to the single selective environment in the simulations. This occurred because the distance threshold used to define connections in the co-association modules was chosen as one that enriched for connections among selected loci with non-random associations in allele frequencies due to selection by a common environmental factor (Supplementary Figure 13).

The propensity of neutral loci to form tightly-clustered co-association networks increased with the complexity of demographic history (compare Figure 8 IBD in left column to 2R in right column). For example, the false positive neutral loci from the two refugia (2R) model formed tightly connected networks, despite the fact that all simulated loci were unlinked. This occurred because of non-random associations in allele frequency due to a shared demographic history.

In some cases, selected loci formed separate or semi-separate modules according to their strengths of selection, but the underlying patterns of association were the same (e.g. Figure 8A, Supplementary Figure 14).

**Figure 8.**
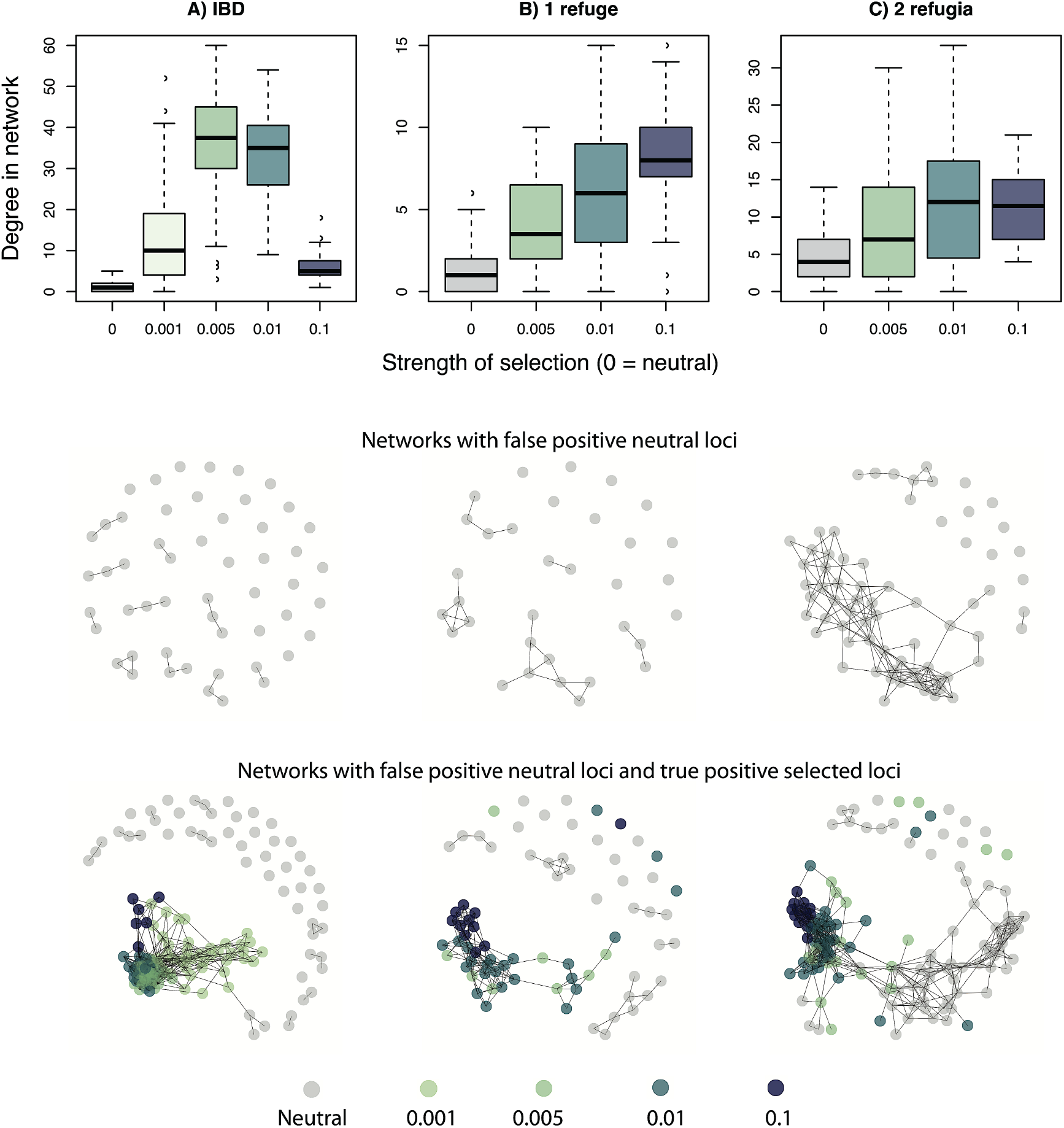
Comparison of co-association networks resulting from simulated data for 3 de-mogra-phies. A) Isolation by distance (IBD), B) range expansion from a single refuge (1R), and C) range expansion from two refugia (2R). All SNPs were simulated unlinked and 1% of SNPs were simulated under selection to an unmeasured weak latitudinal cline. Boxplots of degree of connectedness of a SNP as a function of its strength of selection, across all replicate simulations (top row). Examples of networks formed by datasets that were neutral-only (middle row) or neutral+selected (bottom row) outlier loci.

## Discussion

Co-association networks provide a valuable framework for interpreting the genetic architecture of local adaptation to the environment in lodgepole pine. Our most interesting result was the discovery of low recombination rates among genes putatively adapting to different and distinct aspects of climate, which was unexpected because selection is predicted to increase recombination between loci acted on by different sources of selection as discussed below. If the loci we studied were true causal loci, then different sources of selection were strong enough to reduce LD among *physically linked* loci in the genome, resulting in modular effects of loci on fitness in the environment. While the top candidate SNPs from most genes had associations with only a single environmental factor, for some genes we discovered evidence of environmental pleiotropy, i.e., candidate SNPs associated with multiple distinct aspects of climate. Within co-association modules, we observed a combination of local sweep-like signatures (in which derived alleles at a locus were all found in a particular climate, e.g., cold environments) and antagonistically pleiotropic patterns underlying adaptation to climate (in which some derived alleles at a locus were found at one environmental extreme and others found at the opposite extreme), although we could not evaluate the relative importance of these patterns. Finally, we observed that the modularity of candidate genes in their transcriptionally plastic responses to climate factors did not correspond to the modularity of these genes in their patterns of association with climate, as evidenced through comparing co-association networks with co-expression networks. These results give insight into evolutionary debates about the extent of modularity and pleiotropy in the evolution of genetic architecture [18–24].

### Genetic architecture of adaptation: pleiotropy and modularity

Most of the top candidate genes in our analysis do not exhibit universal pleiotropy to distinct aspects of climate as defined by the expected pattern outlined in Figure 1B. Our results are more consistent with the the Hypothesis of Modular Pleiotropy [19], in which loci may have extensive effects *within* a distinct aspect of the environment (as defined by the variables that associate with each co-association module), but few pleiotropic effects *among* distinct aspects of the environment. These results are in line with theoretical predictions that modular architectures should be favored when there are many sources of selection in complex environments [26]. But note also that if many pleiotropic effects are weak, the stringent statistical thresholds used in our study to reduce false positives may also reduce the extent to which pleiotropy is inferred [20, 21]. Therefore in our study, any pleiotropic effects of genes on fitness detected in multiple aspects of climate are likely to be large effects, and we refrain to making any claims as to the extent of environmental pleiotropy across the entire genome.

The extent of pleiotropy *within* individual co-association modules is hard to quantify, as for any given module we observed associations between genes and several environmental variables. Associations between a SNP and multiple environmental variables may or may not be interpreted as extensive environmental pleiotropic effects, depending on whether univariate environmental variables are considered distinct climatic factors or collectively represent a single multivariate optimum. In many cases, these patterns are certainly affected by correlations among the environmental variables themselves.

Our results also highlight conceptual issues with the definition of and interpretation of pleiotropic effects on distinct aspects of fitness from real data: namely, what constitutes a “distinct aspect” (be it among traits, components of fitness, or aspects of the environment)? In this study we defined the selective environment through the perspective of those environmental variables we tested for associations with SNPs, using a threshold that produced reasonable results in simulation. But even with this definition, some co-association modules are more similar in their multivariate environmental “niche” than others. For instance, genes within the Geography group could be interpreted to have extensive pleiotropic effects if the patterns of associations of each individual module were taken to be “distinct,” or they may be considered to have less extensive pleiotropic effects if their patterns of associations were too similar to be considered “distinct.” While the framework we present here is a step toward understanding and visualizing this hierarchical nature of “distinct aspects” of environmental factors, a more formal framework is needed to quantify the distinctness of pleiotropic effects.

### Genetic architecture of adaptation: linkage

We also observed physical linkage among genes that were associated with very distinct aspects of climate. This was somewhat unexpected from a theoretical perspective: while selection pressures due to genome organization may be weak, if anything, selection would be expected to disfavour linkage and increase recombination between genes adapting to selection pressures with different spatial patterns of variation [34–36]. Interestingly, while the linkage map suggests that these loci are sometimes located relatively close together on a single chromosome, this does not seem to be sufficient physical linkage to also cause a noticeable increase in LD. In other words, it is possible that the amount of physical linkage sometimes observed between genes in different co-association modules is not strong enough to constrain adaptation to these differing gradients. Genetic maps and reference genomes are not yet well developed for the large genomes of conifers; improved genetic maps or assembled genomes will be required to explore these questions in greater depth. If this finding is robust and not compromised by false positives, physical linkage among genes adapting to different climatic factors could either facilitate or hinder a rapid evolutionary response as the multivariate environment changes [4, 5].

Within co-association modules, we observed varying patterns of physical linkage among genes. The Aridity group, in particular, consisted of several tightly linked genes that may have arisen for a number of different reasons. Clusters of physically linked genes such as this may act as a single large-effect QTL [64] and may have evolved due to competition among alleles or genomic rearrangements [30, although these are rare in conifers], increased establishment probability due to linked adaptive alleles [4], or divergence within inversions [32]. Alternatively, if the Aridity region was one of low recombination, a single causal variant could create the appearance of linked selection [65], a widespread false positive signal may have arisen due to genomic variation such as background selection and increased drift [66–68], or a widespread false signal may have arisen due to a demographic process such as allele surfing [69, 70].

### Genetic architecture of adaptation: modularity of transcriptional plasticity vs. fitness

We also compared co-expression networks to co-association networks. Genes that showed similar responses in expression in lodgepole pine seedlings in response to experimental climatic treatments form a co-expression network. Since co-expression networks have been successful at identifying genes that respond the same way to environmental stimuli [71], it might be reasonable to expect that if these genes were adapting to climate they would also show similar patterns of associations with climate variables. However, differential expression analyses only identify genes with plastic transcriptional responses to climate. Plasticity is not a prerequisite for adaptation and may be an alternative strategy to adaptation. This is illustrated by our result that only half of our top candidate contigs for adaptation to climate were differentially expressed in response to climate conditions.

Interestingly, loci located within the same co-association module (groups of loci that are putatively favored or linked to loci putatively favored by natural selection) could be found in different co-expression clusters. For example, we observed that loci from the tightly linked Aridity module had many distinct expression patterns in response to climate treatments. Conversely, candidate genes that were associated with different aspects of the multivariate environment (because they were located in different co-association modules) could nonetheless be co-expressed in response to specific conditions. These observations support the speculation that the developmental/functional modularity of plasticity may not correspond to the modularity of the genotype to fitness map; however, the power of the analysis could be low due to stringent statistical cutoffs and these patterns warrant further investigation.

### Physiological adaptation of lodgepole pine to climate

It is challenging to disentangle the physiological effects and importance of freezing versus drought in the local adaptation of conifers to climate. We found distinct groups of candidate genes along an axis of warm/wet to cold/dry (co-association modules in the Freezing and Multi groups), and another distinct group along an axis of cold/wet to warm/dry (the Aridity co-association module). Selection by drought conditions in winter may occur through extensive physiological remodeling that allows cells to survive intercellular freezing by desiccating protoplasts - but also results in drought stress at the cellular level [55]. Another type of winter drought injury in lodgepole pine - red belt syndrome - is caused by warm, often windy events in winter, when foliage desiccates but the ground is too cold for roots to be able to supply water above ground [72]. This may contrast with drought selection in summer, when available soil water is lowest and aridity highest. The physiological and cellular mechanisms of drought and freezing response have similarities but also potentially important differences that could be responsible for the patterns we have observed.

Our results provide a framework for developing hypotheses that will help to disentangle selective environments and provide genotypes for assisted gene flow in reforestation [73]. While climate change is expected to increase average temperatures across this region, some areas are experiencing more precipitation than historic levels and others experiencing less [74]. Tree mortality rates are increasing across North America due to increased drought and vapour pressure deficit for tree species including lodgepole pine, and associated increased vulnerability to damaging insects, but growth rates are also increasing with warming temperatures and increased carbon dioxide [75, 76]. Hot, dry valleys in southern BC are projected to have novel climates emerge that have no existing analogues in North America [77]. The considerable standing adaptive variation we observe here involving many genes could facilitate adaptation to new temperature and moisture regimes, or could hinder adaptation if novel climates are at odds with the physical linkage among alleles adapted to different climate stressors.

### Limitations of associations with principal components

For these data, testing associations of genes with PC-based climate variables would have led to a very limited interpretation of the environmental drivers of selection because the PC ordination is not biologically informed as to what factors are driving divergent selection [37]. First, many putative candidates in the Freezing and Geography groups would have been missed. Second, strong associations between the Multi SNPs and environmental variables that did not load strongly onto PC1, such as latitude, would have also been missed. Finally, many Aridity SNPs were outliers in PC3, which was strongly correlated with variables that the Aridity SNPs did not have any significant associations with. This occurred because no single variable loaded strongly onto PC3 (the maximum loading of any single variable was 0.38) and many variables had moderate loadings, such that no single environmental variable explained the majority of the variance (the maximum variance explained by any one variable was 15%). Thus, associations with higher PC axes become increasingly difficult to interpret when the axis itself explains less variance of the multivariate environment and the environmental factors loading onto that axis explain similar amounts of variance in that axis. While principal components will capture the environmental factors that covary the most, this may have nothing to do with the combinations that drive divergent selection and local adaptation. This needlessly adds a layer of complexity to an analysis that may not reveal anything biologically important. In contrast, co-association networks highlight those combinations of environments that are biologically important for those genes likely involved in local adaptation.

### Benefits and caveats of co-association networks

Co-association networks provide an intuitive and visual framework for understanding patterns of associations of genes and SNPs across many potentially correlated environmental variables. By parsing loci into different groups based on their associations with multiple variables, this framework offers a more informative approach than grouping loci according to their outlier status based on associations with single environmental variables. While in this study we have used them to infer groups of loci that adapt to distinct aspects of the multivariate environment, co-association networks could be widely applied to a variety of situations, including genotype-phenotype associations. They offer the benefit of jointly identifying modules of loci and the groups of environmental variables that the modules are associated with. While the field may still have some disagreement about how modularity and pleiotropy should be defined, measured, and interpreted [19–21, 23, 24], co-association networks at least provide a quantitative framework to define and visualize modularity.

Co-association networks differ from the application of bipartite network theory for estimating the degree of classical pleiotropic effects of genes on traits [3]. Bipartite networks are two-level networks where the genes form one type of nodes and the traits form the second type of nodes, then a connection is drawn from a gene to a trait if there is a significant association [3]. The degree of pleiotropy of a locus is then inferred by the number of traits that gene is connected to. With the bipartite network approach, trait nodes are defined by those traits measured, and not necessarily the multivariate effects from the perspective of the gene (e.g., a gene that affects organism size will have effects on height, weight and several other variables - if all these traits are analyzed, this gene would be inferred to have large pleiotropic effects). Even if highly correlated traits are removed, simulations have shown that even mild correlations in mutational effects can bias estimates of pleiotropy from bipartite networks [20, 21]. The advantage of co-association networks is their ability to identify combinations of variables (be they traits or environments) that associate with genetic (or SNP) modules. Correlated variables that measure essentially the same environment or phenotype will simply cluster together in a module, which can facilitate interpretation. On the other hand, correlated variables that measure different aspects of the environment or phenotype may cluster into different modules (as we observed in this study). The observed combinations of associations can then be used to develop and test hypotheses as to whether the genotype-environment combination represents a single multivariate environment that the gene is adapting to (in the case of allele associations with environment or fitness) or a single multivariate trait that the gene affects (in the case of allele associations with phenotypes).

While co-association networks hold promise for elucidating the modularity and pleiotropy of the genotype-phenotype-fitness map, some caveats should be noted. First, correlations among variables will make it difficult to infer the exact conditions that select for or the exact traits that associate with particular allelic combinations. Results from this framework can make it easier, however, to generate hypotheses that can be tested with future experiments. Second, the analysis of simulated data shows that investigators should consider demographic history and choose candidates with caution for data analysis to exclude false positives, as we have attempted here. Co-association networks can arise among unlinked neutral loci by chance, and it is almost certain that some proportion of the “top candidate SNPs” in this study are false positives due to linkage with causal SNPs or due to demographic history. The simulated data also showed, however, that causal SNPs tend to have a higher degree of connection in their co-association network than neutral loci, and this might help to prioritize SNPs for follow up experiments, SNP arrays, and genome editing. Third, it may be difficult to draw conclusions about the level of modularity of the genetic architecture. The number of modules may be sensitive to the statistical thresholds used to identify top candidate SNPs [20, 21] as well as the distance threshold used to identify modules. With our data, the number of co-associations modules and the number of SNPs per module were not very sensitive to increasing this threshold by 0.05, but our results were sensitive to decreasing the threshold 0.05 (a stricter threshold resulted in smaller modules of SNPs with extremely similar associations, and a large number of “modules” comprised of a single SNP unconnected to other SNPs, even SNPs in the same gene) (results not shown). While inferred modules composed of a single SNP could be interpreted as unique, our simulations also show that neutral loci are more likely to be unconnected in co-association networks. Many alleles of small effect may be just below statistical detection thresholds, and whether or not these alleles are included could profoundly change inference as to the extent of pleiotropy [20, 21]. This presents a conundrum common to most population genomic approaches for detecting selection, because lowering statistical thresholds will almost certainly increase the number of false positives, while only using very stringent statistical thresholds may decrease the probability of observing pleiotropy if many pleiotropic effects are weak [20]. Thus, while co-association networks are useful for identifying SNP modules associated with correlated variables, further work is necessary to expand this framework to quantitatively measure pleiotropic effects in genomes.

## Conclusions

In this study we discovered physical linkage among loci putatively adapting to different aspects of the climate. These results give rare insight into both the ecological pressures that favor the evolution of modules by natural selection [19] and into the organization of genetic architecture itself. As climate changes, the evolutionary response will be determined by the extent of physical linkage among these loci, in combination with the strength of selection and phenotypic optima across environmental gradients, the scale and pattern of environmental variation, and the details of migration and demographic fluctuations across the landscape. While theory has made strides to provide a framework for predicting the genetic architecture of local adaptation under divergence with gene flow to a single environment [4, 30, 31, 78–82]. as well as the evolution of correlated traits under different directions and/or strengths of selection when those traits have a common genetic basis [35, 36], how genetic architectures evolve on complex heterogeneous landscapes has not been clearly elucidated. Furthermore, it has been difficult to test theory because the field still lacks frameworks for evaluating empirical observations of adaptation in many dimensions. Here, we have attempted to develop an initial framework for understanding adaptation to several complex environments with different spatial patterns, which may also be useful for understanding the genetic basis of multivariate phenotypes from genome-wide association studies. This framework lays the foundation for future studies to study modularity across the genotype-phenotype-fitness continuum.

## Methods

### Sampling and climate

This study uses the same dataset analyzed by Yeaman et al. [46], but with a different focus as explained in the introduction. Briefly, we obtained seeds from 281 sampling locations of lodgepole pine from reforestation collections for natural populations, and these locations were selected to represent the full range of climatic and ecological conditions within the species range in British Columbia and Alberta based on ecosystem delineations. Seeds were grown in a common garden and 2–4 individuals were sampled from each sampling location. The environment for each sampling location was was characterized by estimating climate normals for 1961–1990 from geographic coordinates using the software package ClimateWNA [83]. The program extracts and downscales the moderate spatial resolution generated by PRISM [84] to scale-free and calculates many climate variables for specific locations based on latitude, longitude and elevation. The downscaling is achieved through a combination of bilinear interpolation and dynamic local elevational adjustment. We obtained 19 climatic and 3 geographical variables (latitude, longitude, and elevation). Geographic variables may correlate with some unmeasured environmental variables that present selective pressure to populations (e.g., latitude correlates with day length). Many of these variables were correlated with each other on the landscape (Figure 2A).

### Sequencing, bioinformatics, and annotation

The methods for this section are identical to those reported in [46]. Briefly, DNA from frozen needle tissue was purified using a Macherey-Nagel Nucleospin 96 Plant II Core kit automated on an Eppendorf EpMotion 5075 liquid handling platform. One microgram of DNA from each individual tree was made into a barcoded library with a 350 bp insert size using the BioO NEXTflex Pre-Capture Combo kit. Six individually barcoded libraries were pooled together in equal amounts before sequence capture. The capture was performed using custom Nimblegen SeqCap probes [46 for more details, see 47] and the resulting captured fragments were amplified using the protocol and reagents from the NEXTflex kit. All sample preparation steps followed the recommended protocols provided. After capture, each pool of six libraries was combined with another completed capture pool and the 12 individually barcoded samples were then sequenced, 100 base pair paired-end, on one lane of an Illumina HiSeq 2500 (at the McGill University and Genome Quebec Innovation Centre).

Sequenced reads were filtered and aligned to the loblolly pine genome [85] using bwa mem [86] and variants were called using GATK Unified Genotyper [87], with steps included for removal of PCR duplicates, realignment around indels, and base quality score recalibration [46, 87]. SNPs calls were filtered to eliminate variants that did not meet the following cutoffs: quality score >=20, map quality score >= 45, FisherStrand score <= 33, HaplotypeScore <= 7, MQRankSumTest <= −12.5, ReadPosRankSum >−8, and allele balance < 2.2, minor allele frequency > 5%, and genotyped successfully in >10% of individuals. Ancestral alleles were coded as a 0 and derived alleles coded as a 1 for data analysis.

We used the annotations developed for pine in [46]. Briefly, we performed a BLASTX search against the TAIR 10 protein database and identified the top blast hit for each transcript contig (e-value cut-off was 10^−6^). We also performed a BLASTX against the nr database screened for green plants and used Blast2GO [88] to assign GO terms and enzyme codes [46 for details, see 55]. We also assigned GO terms to each contig based on the GO *A. thaliana* mappings and removed redundant GO terms. To identify if genes with particular molecular function and biological processes were over-represented in top candidate genes, we performed a GO enrichment analysis using topGO [89]. All GO terms associated with at least two candidate genes were analyzed for significant over-representation within each group and in all candidate genes (FDR 5%).

### Top Candidate SNPs

First, top candidate genes were obtained from [46]. For this study, genes with unusually strong signatures of association from multiple association tests (uncorrected genotype-phenotype and genotype-environment correlations, for details see [46]) were identified as those with more outlier SNPs than expected by random with a probability of *P* < 10^−9^, which is a very restrictive cutoff (note that due to non-independence among SNPs in the same contig, this *P*-value is an index, and not an exact probability). Thus, the subsequent analysis is limited to loci that we have the highest confidence are associated with adaptation as evidenced by a large number of significant SNPs (not necessarily the loci with the largest effect sizes).

For this study, we identified top candidate SNPs within the set of top candidate genes. These “top candidate SNPs” had genetic-environment associations with (i) *P*-values lower than the Bonferroni cutoff for the uncorrected Spearman’s ρ (~10^-8^ = 0.05/(number of SNPs times the number of environmental variables) and (ii) log_10_(BF) > 2 for the structure-corrected Spearman’s ρ (Bayenv2, for details see below). The resulting set of candidate SNPs reject the null hypothesis of no association with the environment with high confidence. In subsequent analyses we interpret the results both before and after correction for population structure, to ensure that structure correction does not change our overall conclusions. Note that because candidate SNPs are limited to the top candidate genes in order to reduce false positives in the analysis, these restrictive cutoffs may miss many true positives.

For uncorrected associations between allele frequencies and environments, we calculated the non-parametric rank correlation Spearman’s ρ between allele frequency for each SNP and each environmental variable. For structure-corrected associations between allele frequencies and environments, we used the program Bayenv2 [39]. Bayenv2 is implemented in two steps. In the first step the variance-covariance matrix is calculated from allelic data. As detailed in [46]set of non-coding SNPs to calculated the variance-covariance matrix from the final run of the MCMC after 100,000 iterations, with the final matrix averaged over 3 MCMC runs. In the second step, the variance-covariance matrix is used to control for evolutionary history in the calculation of test statistics for each SNP. For each SNP, Bayenv2 outputs a Bayes factor (a value that measures the strength of evidence in favor of a linear relationship between allele frequencies and the environment after population structure is controlled for) and Spearman’s ρ (the non-parametric correlation between allele frequencies and environment variables after population structure is controlled for). Previous authors have found that the stability of Bayes factors is sensitive to the number of iterations in the MCMC [90]. We ran 3 replicate chains of the MCMC with 50,000 iterations, which we found produced stable results. Bayes factors and structure-corrected Spearman’s ρ were averaged over these 35 replicate chains and these values were used for analysis.

### Co-association networks

We first organized the associations into a matrix with SNPs in columns, environments in rows, and the specific SNP-environment association in each cell. These data were used to calculate pairwise Euclidean distances between SNPs based on their associations, and this distance matrix was used to cluster SNP loci with Ward’s hierarchical clustering using the hclust package in R. As described in the results, this resulted in 4 main groups in the data. For each of these main groups, we used undirected graph networks to visualize submodules of SNPs. Nodes (SNPs) were connected by edges if they had a pairwise Euclidean distance less than 0.1 from the distance matrix described above. We found that the results were not very sensitive to this distance threshold. Co-association networks were visualized using the igraph package in R v 1.0.1 [91].

### Linkage disequilibrium

Linkage disequilibrium was calculated among pairwise combinations of SNPs within genes (genes). Mean values of Pearson’s correlation coefficient squared (*r^2^*) were estimated across all SNPs annotated to each pair of individual genes, excluding SNPs genotyped in fewer than 250 individuals (to minimize the contribution of small sample sizes to the calculation of gene-level means).

### Recombination rates

An Affymetrix SNP array was used to genotype 95 full-sib offspring from a single cross of two parents. Individuals with genotype posterior probabilities of > 0.001 were filtered out. This array yielded data for 13,544 SNPs with mapping-informative genotypes. We used the package “onemap” in R with default settings to estimate recombination rates among pairs of loci, retaining all estimates with LOD scores > 3 [92]. This dataset contained 2760 pairs of SNPs that were found together on the same genomic contig, separated by a maximum distance of 13k base pairs. Of these 7,617,600 possible pairs, 521 were found to have unrealistically high inferred rates of recombination (r > 0.001), and are likely errors. These errors probably occurred as a result of the combined effect of undetected errors in genotype calling, unresolved paralogy in the reference genome that complicates mapping, and differences between the reference loblolly genome that was used for SNP design and the lodgepole pine genomes. As a result, recombination rates that were low (r < 0.001) were expected to be relatively accurate, but we do not draw any inferences about high recombination estimates among loci.

### Associations with principal components of environments

To compare inference from co-association networks to another multivariate approach, we conducted a principal components analysis of environments using the function prcomp() in R. Then, we used Bayenv2 to test associations with PC axes as described above and used BF > 2 as criteria for significance of a SNP on a PC axis. Note that this criterion is less conservative than that used to identify candidate SNPs for the network analysis (because it did not require the additional criteria of a significant Bonferroni-corrected *P*-value), so it should result in greater overlap between PC candidate SNPs and top candidate SNPs based on univariate associations.

### Enrichment of co-expressed genes

The co-expression data used in this study was previously published by [55]. To determine if adaptation cluster members had similar gene functions, we examined their gene expression patterns in response to seven growth chamber climate treatments using previously published RNAseq data [55]. Expression data was collected on 44 seedlings from a single sampling location, raised under common conditions, and then exposed to growth chamber environments that varied in their temperature, moisture and photoperiod regimes. We used a Fisher’s exact test to determine if genes with a significant climate treatment effect were over-represented in each of the 4 major groups and across all adaptation candidates relative to the other sequenced and expressed genes. In addition, Yeaman et al 2014 used weighted gene co-expression network analysis (WGCNA) to identify eight clusters of co-regulated genes among the seven climate treatments. We used a Fisher’s exact test to determine if these previously identified expression clusters were over-represented in the any of the 4 major groups relative to the other sequenced and expressed genes.

### Galaxy biplots

To give insight into how the species has evolved to inhabit multivariate environments relative to the ancestral state, we visualized the magnitude and direction of associations between the derived allele frequency and environmental variables. Allelic correlations with any pair of environmental variables can be visualized by plotting the value of the non-parametric rank correlation Spearman’s *ρ* of the focal allele with variable 1 against the value with variable 2. Spearman’s *ρ* can be calculated with or without correction for population structure. Note also that the specific location of any particular allele in a galaxy biplot depends on the way alleles are coded. SNP data were coded as 0, 1, or 2 copies of the loblolly reference allele. If the reference allele has positive Spearman’s *ρ* with temperature and precipitation, then the alternate allele has a negative Spearman’s *ρ* with temperature and precipitation. For this reason, the alternate allele at a SNP should be interpreted as a reflection through the origin (such that Quadrants 1 and 3 are symmetrical and Quadrants 2 and 4 are symmetrical if the reference allele is randomly chosen).

A prediction ellipse was used to visualize the genome-wide pattern of covariance in allelic effects on a galaxy biplot. For two variables, the 2×2 variance-covariance matrix of *Cov*(*ρ*(*f*, *E*_1_)*, ρ(f, E*_2_)), where *f* is the allele frequency and *E_x_* is the environmental variable, has a geometric interpretation that can be used to visualize covariance in allelic effects with ellipses. The covariance matrix defines both the spread (variance) and the orientation (covariance) of the ellipse, while the expected values or averages of each variable (E[*E*_1_] and E[*E*_2_) represent the centroid or location of the ellipse in multivariate space. The geometry of the two-dimensional (1 − α) × 100% prediction ellipse on the multivariate normal distribution can then be approximated by:

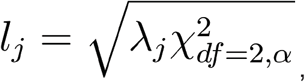
where I*_j_*= {1, 2} represents the lengths of the major and minor axes on the ellipse, respectively, *λ_j_* represents the eigenvalues of the covariance matrix, and 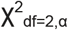 represents the value of the X^2^ distribution for the desired *α* value [93–95]. In the results, we plot the 95% prediction ellipse (*α =*0.05) corresponding to the volume within which 95% of points should fall assuming the data is multivariate normal, using the function `ellipsoidPoints ()` in the R package `cluster`. This approach will work when there is a large number of unlinked SNPs in the set being visualized; if used on a candidate set with a large number of linked SNPs and/or a small candidate set with non-random assignment of alleles (i.e., allele assigned according to a reference), the assumptions of this visualization approach will be violated.

### Visualization of allele frequencies on the landscape

ESRI ArcGIS v10.2.2 was used to visualize candidate SNP frequencies across the landscape. Representative SNPs having the most edges within each sub-network were chosen and plotted against climatic variables representative of those co-association modules. Mean allele frequencies were calculated for each sampled population and plotted using ESRI ArcGIS v10.2.2. Climate data and 1 km resolution rasters were obtained using ClimateWNA v5.40 [83] and shaded with colour gradients scaled to the range of climates across the sampling locations. The climates for each sampling location were also plotted, as some sampling locations were at especially high or low elevations relative to their surrounding landscapes. For clarity, only sampling locations containing at least two sampled individuals were plotted.

### Simulations

The simulations used in this study are identical a subset of those previously published by [62, 63]. Briefly, the simulator uses forward-in-time recurrence equations to model the evolution of independent haploid SNPs on a quasi-continuous square landscape. We modelled three demographic histories that resulted in the same overall neutral *F_ST_* for each demography, but demographic history determined the distribution of *F_ST_’s* around that mean. Isolation by distance (IBD) had the lowest variance, followed by demographic expansion from a single refuge (1R), and demographic expansion from two refugia 2R had the highest variance. The landscape size was 360 × 360 demes and migration was determined by a discretized version of a Gaussian dispersal kernel. Carrying capacity per deme differed slightly for each scenario to give the same overall neutral *F_ST_=* 0.05. IBD was run until equilibrium at 10,000 generations, but 1R and 2R were only run for 1,000 generations in order to mimic the the expansion of lodgepole pine since the last glacial maximum [96]. All selected loci adapted to computer generated landscape with a weak north-south cline and spatial heterogeneity at smaller spatial scales with varying strengths of selection from weak (*s* = 0.001) to strong (*s* = 0.1). See [62, 63] for more details.

The simulations were then expanded in the following way: for each of the 22 environmental variables for lodgepole pine populations, we used interpolation to estimate the value of the variable at the simulated locations. This strategy preserved the correlation structure among the 22 environmental variables. For each of the 22 variables, we calculated the uncorrected rank correlation (Spearman’s *ρ*) between allele frequency and environment. The 23rd computer-generated environment was not included in analysis, as it was meant to represent the hypothetical situation that there is a single unmeasured (and unknown) environmental variable that is the driver of selection. The 23rd environment was correlated from 0–0.2 with the other 22 variables.

We compared two thresholds for determining which loci were retained for co-association network analysis, keeping loci with either: (i) a *P*-value lower than the Bonferroni correction (0.05/(# environments * # simulated loci)) and (ii) a log-10 Bayes Factor greater than 2 (for at least one of the environmental variables). Using both criteria is more stringent and both were used in the lodgepole pine analysis. In the simulations, however, we found that using both criteria resulted in no false positives in the outlier list (see Results); therefore we used only the first of these two criteria so that we could understand how false positives may affect interpretation of the co-association network analysis. For a given set of outliers (e.g., only false positives or false positives and true positives), hierarchical clustering and undirected graph networks were built in the same manner as described for the lodgepole pine data.

List of abbreviations
LD: Linkage disequilibrium
PC: Principal components
SNP: single nucleotide polymorphism

## Declarations

### Ethics approval and consent to participate

Not applicable.

### Consent for publication

Not applicable.

### Availability of data and material

The datasets and analysis code supporting the conclusions of this article are available in the GtiHub repository https://aithub.com/DrK-Lo/Modularitvcode

### Authors’ contributions

KEL conceived of the analysis, conducted analyses, and lead writing of the manuscript. KH and SY did the bioinformatics and various specific analyses. JD created the allele frequency landscape plots. SA led the AdapTree project. All authors contributed to writing of the manuscript.

## Acknowledgements

We thank Sebastian E. Ramos-Onsins, Tanja Pyhäjärvi, an anonymous reviewer and PCI Evol Biol for comments that greatly improved this manuscript. Mike Whitlock provided valuable advice and feedback on various aspects of the research. We thank Jeremy Yoder for organizing the SNP chip data used for calculating the recombination rates. Pia Smets, Connor Fitzpatrick and Sarah Markert assembled and grew genetic materials, and Kristin Nurkowski prepared sequence capture libraries. Tongli Wang and Andreas Hamann selected populations based on climatic distribution of species. Seeds were kindly donated by 63 forest companies and agencies in Alberta and British Columbia (listed at http://adaptree.forestry.ubc.ca/seed-contributors/).

## Supplementary Tables

**Table S1.** Results from GO analysis for all top candidate genes and for each group. The top 5 processes are shown for each category. “P” represents the *P*-value from parent-child Fisher test, while “fdr” represents significance after correction for false discovery rate.

**Table S2.** Top candidate genes and their annotations. For each gene the following information is indicated: the co-association module ID (“group_subMod”), the number of outlier SNPs in each of the four major groups (“Multi”, “Aridity”, “Freezing”, or “Geography”), the Gene ID used in the main paper (“NewContiglDMod”), the color used for plotting (“module_col”), whether or not its homolog shows convergent signals of adaptation with spruce (“is.covergent”), TAIR ID (“tair”), putative gene function (“Annotations”), whether or not the gene was differentially expressed (“diffExp”), and the co-expression cluster (“coexCluster”).

## Supplementary Figures

**Figure S1.** Histogram of *X^T^X* estimated from Bayenv2 for all SNPs (top) and for top candidate SNPs (bottom).

**Figure S2.** Undirected graph network for the Multi group (enlarged version of Figure 2C).

**Figure S3.** Undirected graph network for the Aridity group (enlarged version of Figure 2D).

**Figure S4.** Undirected graph network for the Freezing group (enlarged version of Figure 2E).

**Figure S5.** Undirected graph network for the Geography group (enlarged version of Figure 2F).

**Figure S6.** Heatmap of structure-corrected allele associations with the environment, analogous to Figure 2B in the main paper. Note that although the pattern is very similar, the magnitude of allele correlations is smaller in the structure-corrected data.

**Figure S7.** Linkage disequilibrium heatmap. Mean correlation among allele frequencies between top candidate genes. Genes are ordered the same as Figure 2G in the main paper.

**Figure S8.** Recombination heatmap, clustered by recombination rates. The same data as is shown in Figure 3, except re-clustered by recombination rates to more easily see the patterns of physical linkage.

**Figure S9.** Loadings of environments onto PC axes. The length and direction of each vector represents the scaled loading of that environmental variable onto the PC axis. The color of each vector represents the mean proportion of variance explained by that environment in the two axes plotted.

**Figure S10.** Outliers on PC axes. The distribution of log-10 Bayes Factors for the association between a SNP and a PC axis. Each point is a SNP colored according to its co-association module in Figure 2C-F. Vertical and horizontal lines represent criteria for significance, and the black ovals represent the 95% prediction ellipse. Note that candidate SNPs all had BF > 2 with at least one univariate environmental variable.

**Figure S11.** SNP annotations and genomic features. Proportion of exome SNPs falling into various categories for genomic features compared to in the top candidate list. 3primeFLANK: 3’ flanking region; 3primeUTR: 3’ untranslated region; 5primeFLANK: 5’ flanking region; 5primeUTR: 5’ untranslated region; non-tcontig: not located in a transcriptomic contig (intergenic); nonsyn: non-synonymous substitution; unk-adj: unknown adjacent region; unk-flank: unknown flanking region; UNKNOWN-ORF: unknown open reading frame.

**Figure S12.** Error rates from the simulations given a less stringent criteria (Bonferroni, left) and a more stringent criteria (Bonferroni and Bayes Factors from bayenv2, right). The less stringent criteria was used for the simulations because it had some false positives (A), while the more stringent criteria was used for the empirical data because it didn’t have any false positives (B). The three demographies are isolation by distance (IBD), range expansion from one refuge (1R), and range expansion from two refugia (2R). While using the more stringent criteria resulted in no false positives, it also reduced the number of true positives (compare C and D), with the most severe reduction under isolation by distance.

**Figure S13.** Pairwise distances among loci as a function of selection for simulated data. Evaluation of 0.1 as a distance threshold for creating an co-association module. The three demographies are isolation by distance (IBD), range expansion from one refuge (1R), and range expansion from two refugia (2R). For the simulated data, top candidates were chosen as described in the methods. Multivariate euclidean distance was calculated among the loci based on their associations with environments, and the proportion of pairwise distances above the distance threshold of 0.1 (used for the empirical data) was calculated for each type of comparison. We evaluated four types of pairwise comparisions: neutral loci with each other (“Neut-Neut”), neutral loci with selected loci (“Neut-Sel”), all selected loci with each other (“Sel-Sel”), and only loci under strong selection with each other (s > 0.1, “strongSel-strongSel”). A higher proportion of pairwise distances above the threshold indicates that these loci would be more connected to each other in the co-association network.

**Figure S14.** More examples of networks from simulations. The simulated datasets were nested within randomly generated selective environments, such that different demographic histories were simulated on the same environmental landscape. For this randomly generated environment, loci simulated under stronger selection had a propensity to cluster differently than loci simulated under weaker selection. To be clear, they still show the same patterns of associations, but the absolute value of the associations was just larger for the loci under strong selection and this caused the creation of a second cluster.

